# Evolution of diapause in the African turquoise killifish by remodeling ancient gene regulatory landscape

**DOI:** 10.1101/2021.10.25.465616

**Authors:** Param Priya Singh, G. Adam Reeves, Kévin Contrepois, Mathew Ellenberger, Chi-Kuo Hu, Michael P. Snyder, Anne Brunet

**Affiliations:** Department of Genetics, Stanford University, Stanford, CA, USA; Stanford Cardiovascular Institute, Stanford University, Stanford, CA, USA; Stanford Diabetes Research Center, Stanford University, Stanford, CA, USA; Glenn Laboratories for the Biology of Aging, Stanford University, Stanford, CA, USA

## Abstract

Suspended animation states such as hibernation or diapause allow organisms to survive extreme environments. But the mechanisms underlying the evolution of these extreme survival states are unknown. The African turquoise killifish has evolved diapause as a form of suspended development to survive the complete drought that occurs every year in its habitat. Here we show that many gene duplicates – paralogs – exhibit specialized expression in diapause versus normal development in the African turquoise killifish. Surprisingly, paralogs with specialized expression in diapause are evolutionarily very ancient, and they are also present even in vertebrates that do not exhibit diapause. Profiling the chromatin accessibility landscape among different fish species reveals an evolutionarily recent increase in chromatin accessibility at these very ancient paralogs, suggesting rewiring of their regulatory landscape. The increase in chromatin accessibility in the African turquoise killifish is linked to the presence of new binding sites for transcription factors (e.g., FOXO, REST, and PPAR), due to both de novo mutations and transposable element insertion. Interestingly, accessible chromatin regions in diapause are enriched for lipid metabolism genes. By performing lipidomics in different fish species, we uncover a specific lipid profile in African turquoise killifish embryos in diapause. Notably, select very long-chain fatty acids are high in diapause, suggesting they may be used for long-term survival in this state. Together, our multi-omic analysis indicates that diapause is driven by regulatory innovation of very ancient gene programs that are critical for survival. Our work also suggests a mechanism for how complex adaptations evolve in nature and offers strategies by which a suspended animation program could be reactivated in other species for long-term preservation.

## Introduction

Extremophiles – species that live in extreme environments – have evolved unique adaptations for survival. Understanding how extreme adaptations evolve can reveal new pathways with important ramifications for survival in all organisms. The African turquoise killifish *Nothobranchius furzeri* is an extremophile for embryo survival. This vertebrate species lives in ephemeral ponds in Zimbabwe and Mozambique that completely dry up for ~8 months each year (*1*). To survive this annual drought, the African turquoise killifish has evolved two key adaptations: a rapid life to successfully reproduce during the rainy season and a form of long suspended animation, with embryos entering diapause and subsisting in the mud during the dry season (*2–5*). Diapause embryos survive for months even years – longer than adult life – without any detectable tradeoff for future life (*6*). Remarkably, diapause embryos already have complex organs and tissues, including a developing brain and heart (*6*). Hence, diapause provides a unique form of long-term protection to a complex organism.

Like other suspended animation states (hibernation, torpor), diapause is a multifaceted and active adaptation. Diapause also exists in other vertebrate species, including mammals (e.g., bear, roe deer, mice) (*7*). As diapause is extreme in the African turquoise killifish, this species provides a model to understand the mechanism and evolution of this suspended animation trait in vertebrates. Many genes involved in chromatin remodeling, metabolism and stress resistance are upregulated in killifish (*6, 8, 9*). Yet, how such an extreme and coordinated program evolved in nature is unknown. Using the lens of evolution to understand diapause could uncover new protective mechanisms for long-term survival and offer a framework for the evolution of extreme adaptations in nature.

### Paralogs that specialize for expression in diapause are evolutionarily very ancient

We asked when the genes expressed in diapause originated in evolutionary time. To this end, we focused on paralogs – duplicated copies of genes (*10, 11*). Paralogs are the primary mechanism by which new genes originate and specialize for new functions or states (*12, 13*) (Fig. 1A). Paralogs also allow for a precise timing of the evolutionary origin of specific genes and they could help explain how the killifish genome can support two seemingly antagonistic traits – rapid life vs. suspended animation. Using phylogenetic inference (see Methods), we find that the African turquoise killifish genome contains 20,091 paralog pairs. We used our previously generated RNA-seq datasets of development and diapause in the African turquoise killifish (*6*) to analyze if the expression pattern of paralogs has diverged in diapause versus normal development states. Interestingly, many paralog pairs show opposing expression, with one gene in the paralog pair highly expressed in diapause (‘*diapause-specialized gene’* e.g., the chromatin modifier *EZH1*) and the other gene in the paralog pair highly expressed in development (‘*development-specialized gene’* e.g., the chromatin modifier *EZH2*) (Fig. 1B and fig. S1). Overall, 6,247 paralog pairs show expression specialization in diapause versus development state (Fig. 1C and fig. S1, Data Files S1 and S2).

**Figure 1.**
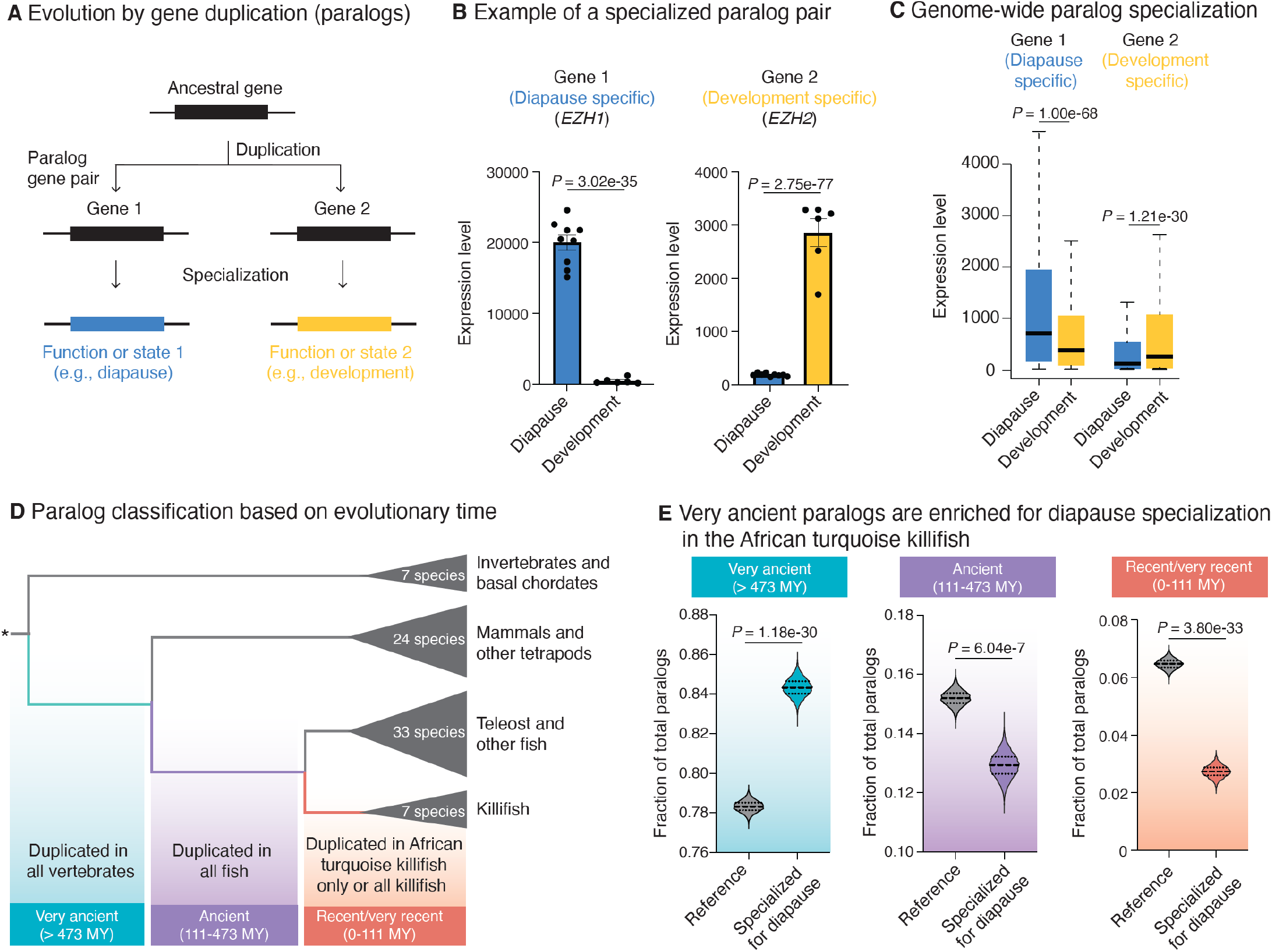
Specialization of very ancient paralogs for expression in diapause in African turquoise killifish. (A) Schematic of paralog specialization after gene duplication. After duplication from the same ancestral gene, genes from a paralog gene pair can specialize for different functions or states (e.g., diapause vs. development). (B) Examples of a paralog gene pair, with specialized expression of gene 1 in diapause (blue, *EZH1*) and gene 2 in development (yellow, *EZH2*) in the African turquoise killifish (*Nothobranchius furzeri*). Bars represent mean expression level (normalized DESeq2 count) across replicates in diapause or development state. Dots show normalized DESeq2 counts in each replicate. Error bar is standard error of mean (SEM). Corrected *P*-values (median from pairwise comparisons) from DESeq2 Wald test. (C) Box plots showing expression levels (normalized DESeq2 counts) of all the specialized paralog pairs in diapause and development in the African turquoise killifish genome. Boxes show the median and interquartile ranges, and whiskers indicate maximum 1.5 interquartile range. Gene 1 of the paralog pair has a higher expression on average in diapause (blue) compared to development (yellow), whereas gene 2 has a higher expression on average in development (yellow) compared to diapause (blue). *P*-values from Kolmogorov-Smirnov test. (D) Schematic for binning paralog duplication time into 3 categories based on OrthoFinder pipeline with 71 species (see Methods and fig. S2 for complete tree). Divergence time estimates are from species tree in Ensembl. Binned categories include genes that were duplicated in the common ancestor of (1) all vertebrates or earlier (very ancient, > 473 million years [MY]), (2) all fish (ancient, 111-473 MY), and (3) all killifish or African turquoise killifish exclusively (recent/very recent, 0-111 MY). (E) Fraction of total paralog pairs within each of the very ancient (left), ancient (middle), and recent/very recent (right) binned categories. Violin plots represent the distribution of observed vs expected specialized paralog fractions generated through 10,000 bootstrapped random sampling. Median and quartiles are indicated by dashed lines. The enrichment of diapause-specialized paralogs pairs within each bin is compared to genome-wide expectation (see Methods). Compared to the reference, paralogs with specialization in diapause are enriched among genes with very ancient duplication times and depleted among genes with ancient and recent/very recent duplication times respectively. *P*-values from Chi-square test.

We next asked whether paralogs that exhibit expression specialization in diapause are evolutionarily recent or ancient. Diapause in the African turquoise killifish is a relatively recent specialization that evolved less than 18 million years (MY) ago (*14*). To date the paralogs, we generated a paralog classification pipeline to identify the evolutionary time when each of the African turquoise killifish paralogs originate compared to other species (Fig. 1D and fig. S2) (*15*). We distinguished i) very ancient paralogs (shared with all vertebrates, including mammals) that originated more than 473 MY ago, ii) ancient paralogs (shared with all other fish) that originated between ~111-473 MY ago, and iii) recent/very recent paralogs (killifish/African turquoise killifish specific) that originated less than ~111 MY ago (Fig. 1D). Surprisingly, very ancient paralogs were significantly more likely to specialize for diapause compared to the genome-wide average, even though diapause originated recently (Fig. 1E). In contrast, ancient and especially recent/very recent (killifish-specific) paralogs were significantly less likely to specialize for diapause compared the genome-wide average, even though they originated around the time when diapause evolved (Fig. 1E). Consistently, paralogs that originate from very ancient vertebrate-specific whole genome duplication or ancient small-scale duplications were more likely to specialize for diapause (fig. S3). The enrichment for very ancient paralog pairs for specialization in diapause was robust to varying outgroups, phylogeny, method to identify paralogs, and paralog family size (fig. S4, A to E). As a specificity control, such an enrichment was not observed for paralogs that are expressed at the same level during development and diapause (in fact, those exhibited a depletion for very ancient paralogs) (fig. S4F). The genes specialized for expression in diapause did not exhibit increased positive selection at the protein level, raising the possibility of regulatory rewiring (fig. S4G). Thus, very ancient paralogs are co-opted in diapause, suggesting that ancestral programs are harnessed for this suspended animation state – perhaps by remodeling of the regulatory landscape.

### Very ancient paralogs also specialize in diapause in other killifish species with diapause

Many killifish species populate the world, and their ability to undergo diapause is linked to their environment. Killifish species that live in ephemeral ponds exhibit diapause (e.g., African turquoise killifish, South American killifish), whereas killifish species that live in constant water do not undergo diapause, and instead they continuously develop (e.g., red-striped killifish and lyretail killifish, both of which are from Africa) (*16–19*) (Fig. 2A). To assess whether the specialization of ancient paralogs in diapause is generalizable to other species that evolved diapause independently, we used available RNA-seq data from diapause and development in the South American killifish with diapause, *Austrofundulus limnaeus* (*9*). We also generated new RNA-seq data from the developing embryos of the red-striped killifish *Aphyosemion striatum* and the lyretail killifish *Aphyosemion australe* – the closest relatives of the African turquoise killifish *N. furzeri* but without diapause (Fig. 2A). We found that in the South American killifish, paralogs also showed specialized expression in diapause versus development (Fig. 2, B and C, and fig. S5), and that their specialized expression correlated with that of paralogs in the African turquoise killifish (Fig. 2D and fig. S6, A and B). In contrast, killifish species without diapause expressed both paralogs during development (fig. S6C). Importantly, paralogs with specific expression in diapause in the South American killifish were also enriched for very ancient gene duplicates (Fig. 2E). Collectively, these results indicate that very ancient paralog pairs have been repeatedly co-opted for specialized expression in diapause during evolution. Together, our observations also raise the possibility that the recent specialization for diapause is driven at least in part by regulatory innovation of very ancient paralog pairs.

**Figure 2.**
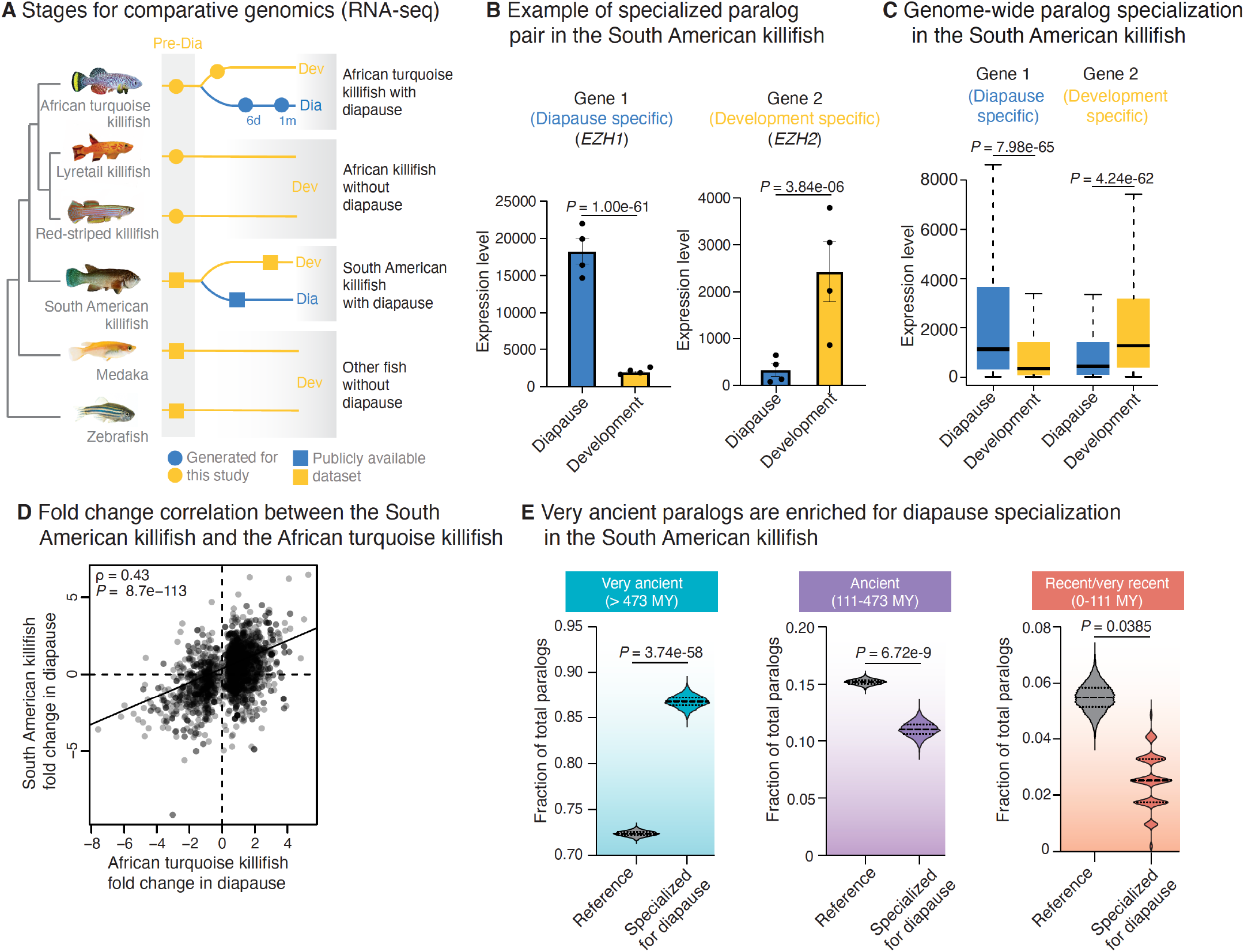
Very ancient paralogs also specialize for expression in diapause in other killifish species with diapause. (A) Experimental design for analysis of RNA-seq datasets either publicly available (square) or *de-novo* generated for this study (circles). Killifish species are from Africa (with and without diapause) and South America (with diapause). Medaka and zebrafish are other teleost fish without diapause. The development stage (yellow square) in the South American killifish corresponds to post-diapause development. Pre-Dia, pre-diapause; Dia, diapause; Dev, development; 6d, 6 days in diapause; 1m, 1 month in diapause. (B) Examples of paralog gene pair, with specialized expression of gene 1 in diapause (blue, *EZH1*) and gene 2 in development (yellow, *EZH2*) in South American killifish (*Austrofundulus limnaeus*). Gene names displayed are the names assigned to the ortholog in African turquoise killifish. Bars represent the mean expression level (normalized DESeq2 count) across replicates in diapause or post-diapause development state. Error bar is standard error of mean (SEM). Each dot represents the normalized expression level for all sample replicates in diapause or post diapause development. *P*-values from DESeq2 Wald test. (C) Box plots showing the expression levels (normalized DESeq2 counts) of all the specialized paralog pairs in diapause and development in South American killifish. Boxes show the median and interquartile ranges, and whiskers indicate maximum 1.5 interquartile range. Gene 1 of the paralog pair has a higher expression on average in diapause (blue) compared to development (yellow), whereas gene 2 has a higher expression on average in development (yellow) compared to diapause (blue). *P*-values from Kolmogorov-Smirnov test. (D) Spearman’s rank correlation between ortholog genes that change with diapause in African turquoise killifish and South American killifish. Dots represent the fold change values of ortholog genes in diapause compared to development in the two species. Spearman’s correlation coefficient (ρ) and *P*-values from t-test are shown on the plot. (E) Fraction of the total paralog pairs within each of the very ancient (left), ancient (middle), and recent/very recent (right) binned categories as described in Fig. 1D. Violin plots represent the distribution of observed vs expected specialized paralog fractions generated through 10,000 bootstrapped random sampling. Median and quartiles are indicated by dashed lines. The enrichment of diapause-specialized paralogs pairs within each bin is compared to genome-wide expectation (reference). Compared to the reference, paralogs with specialization in diapause are enriched among genes with very ancient duplication times and depleted among genes with ancient and recent or very recent duplication times respectively. *P*-values from Chi-square test.

### Evolutionarily recent remodeling of the chromatin landscape at very ancient paralogs

To characterize the regulatory landscape of the paralogs that specialize in diapause during evolution, we profiled the chromatin accessibility landscape in different species of killifish. We performed ATAC-seq (Assay for Transposase-Accessible Chromatin using sequencing), which assesses chromatin accessibility genome-wide (*20*), on embryos during diapause and development in killifish species with diapause (African turquoise killifish, South American killifish) and embryos during development in killifish species without diapause (lyretail killifish and red-striped killifish) at a similar developmental stage (Fig. 3A). We also used available ATAC-seq data for medaka and zebrafish development (*21*). We verified the quality of our ATAC-seq samples by transcription start site enrichment of open chromatin and fragment size periodicity (See Methods, fig. S8). Chromatin states easily separated diapause and development embryos by Principal Component Analysis (PCA) in the African turquoise killifish (Fig. 3B). Chromatin states also separated diapause and developmental samples of different killifish species (Fig. 3B). In the African turquoise killifish, 6,490 genomic regions were differentially accessible in diapause compared to development genome-wide (fig. S7A, Data Files S1 and S3), and they were located mostly in promoter, intronic, or distal intergenic (e.g., enhancer) regions (fig. S7B). There was a positive correlation between chromatin accessibility and gene expression levels in diapause in the African turquoise killifish (fig. S7, C and D). Together, these results indicate that our datasets for chromatin state landscape in diapause and development in several killifish species are of good quality.

**Figure 3.**
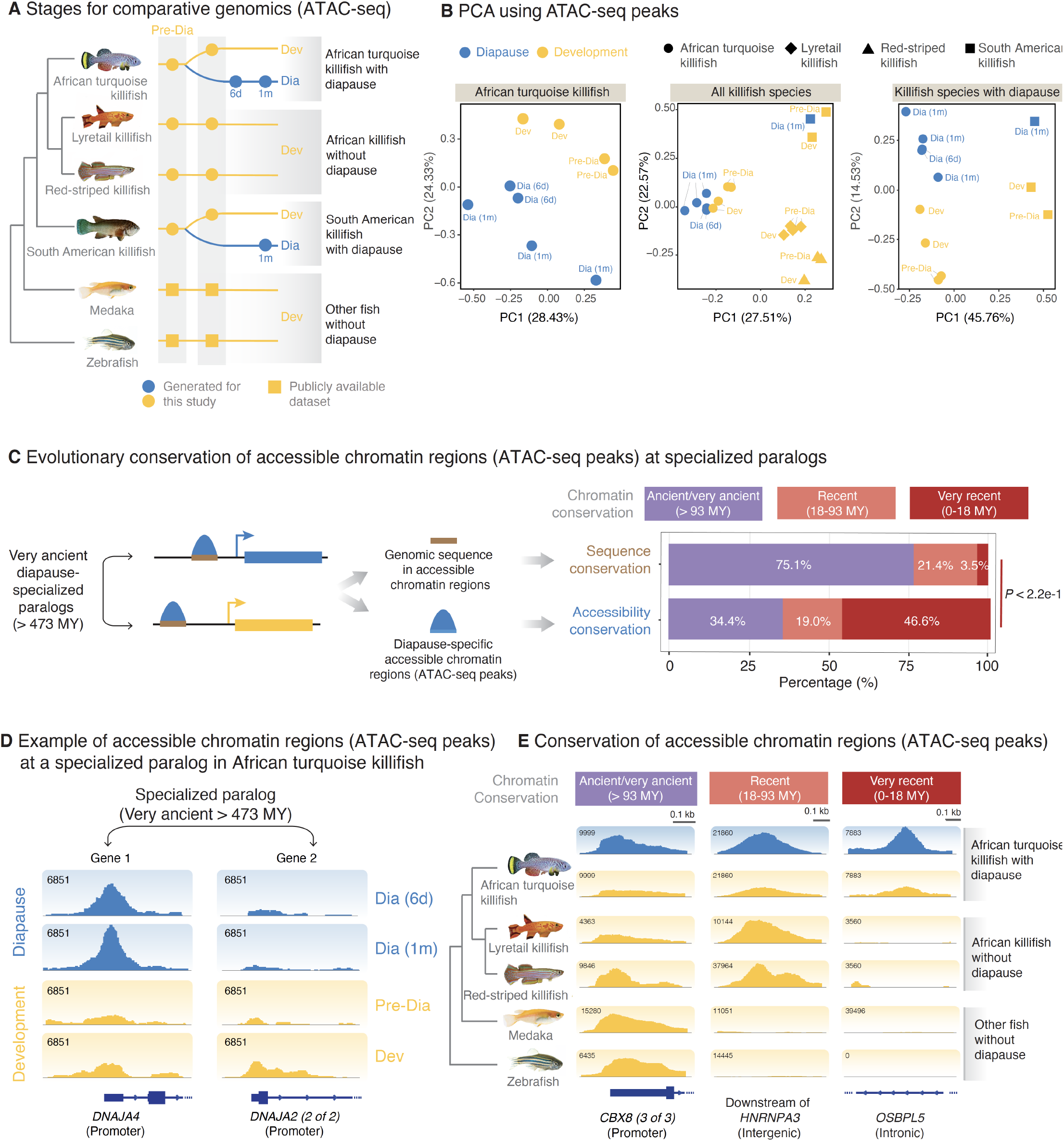
Evolutionarily recent remodeling of genome-wide chromatin landscape drives specialized expression of very ancient paralogs in diapause. (A) Experimental design for the ATAC-seq datasets either publicly available (squares) or de novo generated in this study (circles) (see also Data File S2). Pre-Dia, pre-diapause; Dia, diapause; Dev, development; 6d, 6 days in diapause; 1m, 1 month in diapause. (B) Principal component analysis (PCA) on all chromatin accessibility regions in each species group: African turquoise killifish only (left), all killifish species (center), and diapause-capable killifish (right). Each point represents the consensus ATAC-seq peaks (chromatin accessibility) from an individual replicate of a given species at a given developmental or diapause state. Percentage of the variance explained by each Principal Component (PC) is shown in parentheses. (C) Conservation analysis of genomic sequence and chromatin accessibility at very ancient paralogs with specialization in diapause vs. development. Left: Schematic of the analysis. Right: Percentage (e.g., conservation) of alignable regions containing diapause-specific chromatin accessibility (upper) and the conservation of diapause-specific chromatin accessibility (lower) near specialized ancient paralogs (see also fig. S10). While the majority of genomic sequences are ancient, the chromatin accessibility at those peaks evolved recently in the African turquoise killifish. *P*-values from Chi-square test. (D) Genome browser (IGV) visualization of chromatin accessibility regions (ATAC-seq peaks) at the promoter of genes from a very ancient paralog pair, with one gene specialized for expression in diapause (*DNAJA4*) and the other in development (*DNAJA2*). Replicates within each condition were aggregated by summation for visualization. Blue boxes and lines represent genomic features (exons and introns, respectively). (E) Genome-browser (IGV) visualization of representative diapause-specific chromatin accessibility regions that are: ancient/very ancient (conserved across killifish and medaka or zebrafish; left), recent (conserved in killifish species; middle), or very recent (specific to African turquoise killifish; right). Blue boxes and lines at the bottom represent genomic features for the closest gene (exons and introns, respectively). To generate tracks, RPKM-normalized reads were summed across replicates and biological timepoints (e.g., diapause and development separately) to obtain single tracks for each species.

We next examined accessible chromatin regions (ATAC-seq peaks) at paralogs that are differentially expressed in diapause versus development (Fig. 3C) (e.g., *DNAJA4* and *DNAJA2*, Fig. 3D and fig. S1B). As paralogs that specialize in diapause are very ancient (> 473 MY), we therefore asked when chromatin accessibility occurred in evolutionary time (Fig. 3C and fig. S9). To quantify chromatin accessibility over evolutionary time, we developed a pipeline to identify the relative evolutionary origin of ATAC-seq peak based on multi-genome alignment (see Methods), and classified each ATAC-seq peak as i) ancient/very ancient (i.e. chromatin accessible in all fish species evaluated, such as *CBX8*) (Fig. 3E and fig. S9), ii) recent (i.e. chromatin accessible only in killifish species, such as *HNRNPA3*), and very recent (chromatin accessible only in the African turquoise killifish, such as *OSBPL5)* (Fig. 3E and fig. S9). Interestingly, most regulatory regions of very ancient paralogs (> 473 MY) that are differentially regulated in diapause exhibited chromatin accessibility very recently (~18 MY), only in the African turquoise killifish (Fig. 3C and fig. S10). The very recent chromatin accessibility at very ancient paralogs specialized in diapause was generalizable to non-paralog genes (fig. S10A) and was more pronounced at distal regulatory elements (likely enhancers) (fig. S10B). Thus, the African turquoise killifish exhibits an evolutionary recent remodeling of the chromatin accessibility landscape at very ancient genes.

### Mechanisms underlying the evolution of chromatin accessibility in diapause

What are the mechanisms connecting evolutionary recent chromatin accessibility with diapause? Chromatin regions that opened recently in diapause paralogs in the African turquoise killifish were enriched for transcription factor motifs, including REST, NR2F2, Forkhead transcription factors (e.g., FOXA1, FOXO3), and PPAR (e.g., PPARA) (Fig. 4A). Most diapause-specific transcription factor binding motifs were only enriched in the African turquoise killifish, but not in other closely related fish without diapause (Fig. 4B and fig. S11). Thus, these transcription factor motifs arose very recently in the African turquoise killifish after divergence from other killifish species without diapause and could underlie the evolutionarily recent opening of chromatin at very ancient paralogs.

**Figure 4.**
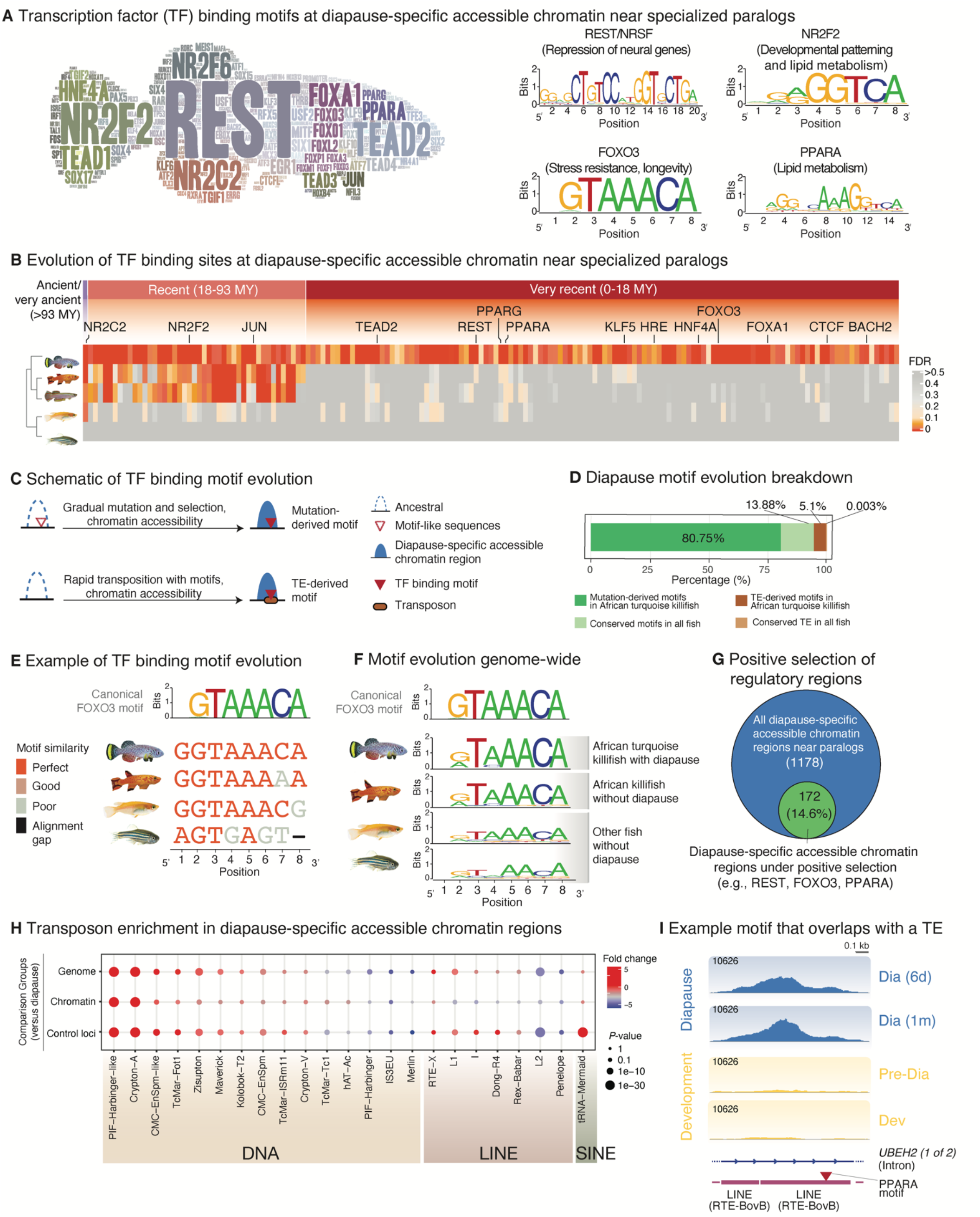
Mechanisms underlying the evolution of chromatin accessibility in diapause at specialized paralogs. (A) Left: Word cloud for transcription-factor binding motifs enriched in the diapause-specific chromatin accessible regions (ATAC-seq peaks) at specialized paralogs using HOMER. Right: Motif logos generated by weblogo for selected transcription factor examples are shown on the right. Y-axis is formatted using *informational content* (i.e., bits) which scales based on single-base overrepresentation in the binding sequence (0: bases are represented equally at 25% each in reference sequences; 2: a single base dominates the entirety of reference sequences at 100%). (B) Conservation in other fish species of transcription-factor binding motifs enriched in the diapause-specific chromatin accessible regions in the African turquoise killifish. The majority of diapause-specific motifs are very recent (i.e., specific to the African turquoise killifish) and not enriched in killifish species without diapause. Selected representative motifs are highlighted (see also figs. S11, S17). (C) Schematic of two possible mechanisms for the evolution of the diapause-specific transcription factor binding motifs in the African turquoise killifish genome. Upper: Gradual mutations paired with selective pressure leads to the formation of binding sites for specific transcription factors and accompanying chromatin accessibility. Lower: A site experiences a transposable element (TE) insertion event, providing a novel sequence that contains a binding site for a specific transcription factor and accompanying chromatin accessibility. (D) Contribution (in percentage) of the two possible evolutionary mechanisms (mutation and TE insertion) for motif evolution in diapause-specific chromatin at specialized paralogs. Diapause-specific motifs likely acquired by mutation in the African turquoise killifish (dark green) are those that are expected to bind their transcription factors exclusively in the African turquoise killifish based on the HOMER log odds ratio binding criteria (see Methods). Conserved motifs likely acquired by mutation in multiple species (light green) are those that are expected to bind their transcription factor motifs in at least one other fish species evaluated. Diapause-specific motifs likely acquired by TE insertion (brown) are those that overlap with an annotated TE sequence and are expected to bind their transcription factor exclusively in the African turquoise killifish. Conserved motifs likely acquired by TE insertion (light brown) are those that overlap with a TE sequence in the African turquoise killifish and at least one other species in addition to being expected to bind their transcription factor in both the African turquoise killifish and at least one additional species. A large majority of motifs likely evolved through mutations in the African turquoise killifish. (E) Example of a transcription factor binding motif that likely evolved via mutation: *FOXO3* binding motif in a diapause-specific chromatin accessible region of the African turquoise killifish genome near *GPC3* (*LOC107379575*) and aligned regions in other fish species. Aligned sequences colored based on the similarity to the canonical *FOXO3* motif from HOMER (top track). (F) Aggregated informational content across all *FOXO3* transcription factor binding sites in diapause-specific accessible chromatin regions and aligned regions in other species. Y-axis is formatted using *informational content* (i.e., bits). The African turquoise killifish motif exhibits greater similarity to the canonical *FOXO3* binding motif (see also fig. S12). (G) Fraction of diapause-specific chromatin accessible regions under positive selection in the African turquoise killifish (see Methods) at FDR = 0.1, which are enriched for selected motifs (e.g., REST, FOXO3, PPARA). (H) Enrichment or depletion of specific transposable elements (TEs) in the diapause-specific chromatin accessible regions (ATAC-seq peaks) in the African turquoise killifish genome as compared to the overall genomic abundance of the given TE (“Genome”), compared to the abundance of the given TE in all chromatin accessible regions (“Chromatin”), and compared to the abundance of the given TE in size-matched control regions 10 kb away from ATAC-seq peak of interest (“Control loci”). (I) Example of a diapause-specific chromatin accessibility region containing a PPARA binding site overlapping with a TE. Almost all TE-derived motifs are specific to the African turquoise killifish.

New transcription binding motifs can arise de novo by point mutation or transposable element (TE) insertion (*22, 23*) (Fig. 4C). A majority (81%) of the transcription factor binding motifs associated with diapause accessible chromatin at specialized paralogs in the African turquoise killifish evolved *de novo*, via mutation of the ancestral sequence (Fig. 4D). For example, transcription factor motifs (e.g., FOXO3 motifs) were canonical binding sites (as defined by HOMER (*24*)) in the African turquoise killifish sequence but were slightly divergent in closely related fish without diapause and even more divergent or even absent in more distant fish species (Fig. 4, E and F, and fig. S12). Importantly, we found a signature of positive selection (*25*) at many of the diapause-specific accessible chromatin regions in the African turquoise killifish, with enrichment for binding motifs for FOXO3, REST, and PPAR (Fig. 4G and fig. S13). Thus, the African turquoise killifish may have selected for canonical transcription factor binding motifs at regulatory regions of genes beneficial for diapause.

Intriguingly, some binding motifs associated with diapause in the African turquoise killifish paralogs overlapped with transposable elements and were unique to this species (Fig. 4D). While overlaps with transposable elements represents a minority of cases (5%) (Fig. 4D), transposable elements can deliver a transcription factor binding motif to new regulatory sites faster than gradual mutation and selection. As transposable elements have exploded in the African turquoise killifish genome (*26*), they may represent an evolutionary mechanism to co-opt genes into the diapause expression program. Several transposable element families (e.g., DNA transposons and LINEs) were highly enriched at accessible chromatin regions in diapause in the African turquoise killifish (Fig. 4H). In some cases, these regions contained both a transposable element and a transcription factor binding site (e.g., PPAR) (Fig. 4I and fig. S14). Hence, transcription factor binding motifs underlying diapause-specialized paralogs may have originated not only through mutation and selection but also via a recent burst, in the African turquoise killifish, of transposon-mediated reshuffling.

### Functional enrichment and lipidomics reveal specific lipids in the diapause state

We asked whether ancient paralogs that are recently repurposed for diapause in the African turquoise killifish are associated with a specific biological function. Analysis of gene expression and chromatin accessibility at diapause-specific paralogs showed enrichment of several functions related to lipid metabolism (e.g., lipid storage, very long chain fatty acid metabolism and regulation of fatty acid beta oxidation) (Fig. 5A, Data Files S5 and S6). Both gene expression and chromatin accessibility datasets showed enrichment of upstream regulators of lipid metabolism (e.g., FOXO1 (*27*)) or transcriptional sensors of fatty acids (e.g., PPAR) (*28*) (Fig. 5B, Data File S7). These observations raise the possibility that lipids play an important role in diapause.

**Figure 5.**
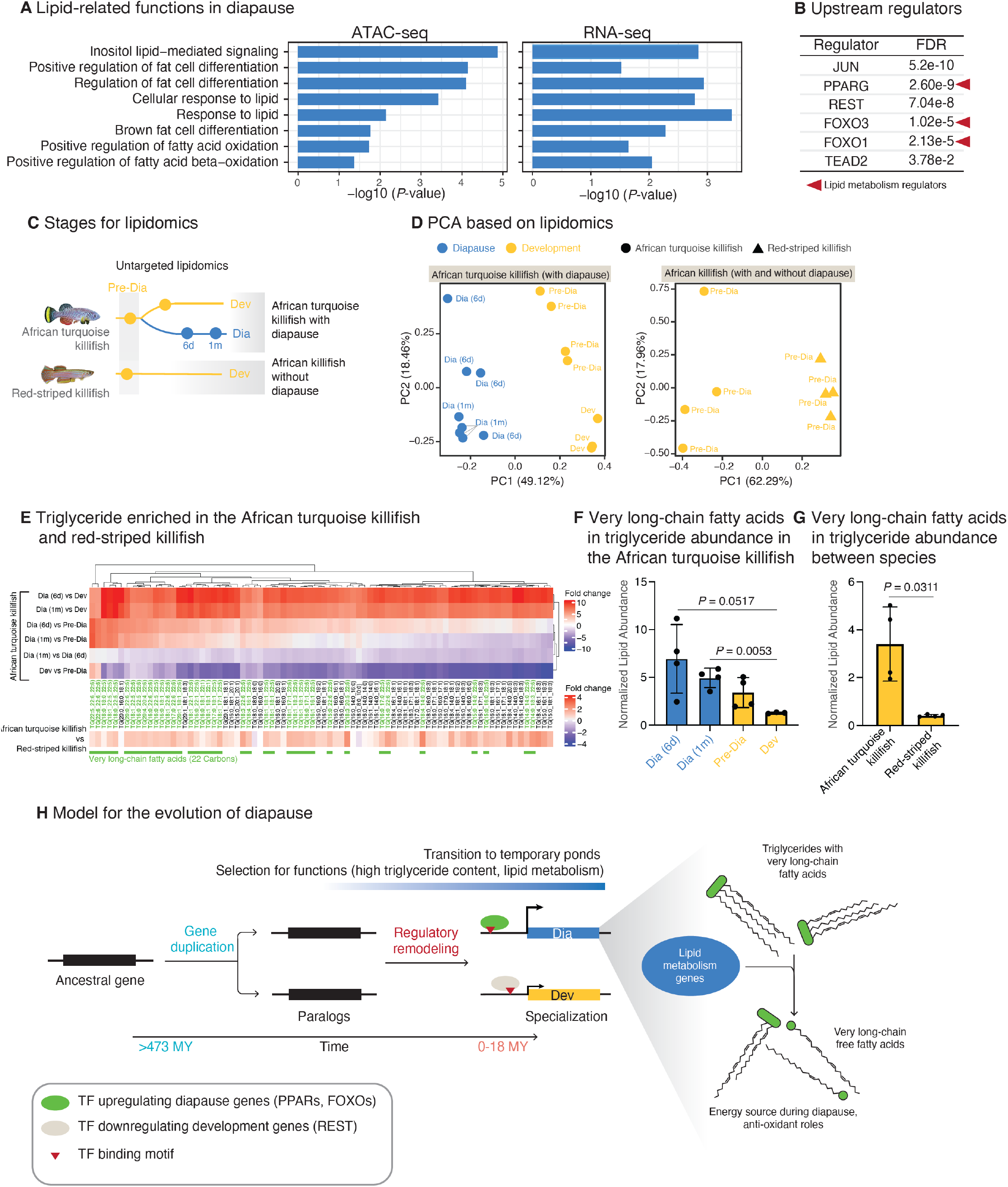
Functional enrichment and lipidomics reveal specific lipids in the diapause state. (A) Gene Ontology enrichment for the lipid-related terms enriched in both expressed genes (diapause specialized paralogs) and chromatin accessible regions in diapause (see Data File S4 for full list). (B) Selected upstream regulators of the paralog genes specialized in diapause using Ingenuity Pathway Analysis (see Data File S4 for full list). Regulators related to lipid metabolism are highlighted (red arrows). Several upstream regulatory transcription factors in this analysis overlap with transcription factor binding motifs enriched in diapause-specific chromatin accessible regions (see Fig. 3). (C) Experimental design for untargeted lipidomics in the African turquoise killifish (with diapause) and the red-striped killifish (without diapause). The two development and diapause time points in the African turquoise and pre-diapause in red-striped killifish are identical to those for ATAC-seq (Fig. 3A). (D) Principal component analysis (PCA) on the estimated concentrations for all detected lipids for the African turquoise killifish only (left) and for both killifish (right). Each point represents an individual replicate from a given species at a given developmental or diapause stage. Variance explained by each Principal Component (PC) is shown in parentheses. (E) Heatmap representing the fold change of significant triglycerides between diapause vs. development in the African turquoise killifish (upper) and between the African turquoise killifish vs. red-striped killifish (development only, lower). Fold change values are plotted between each pair-wise comparison between diapause and development time points, or the two development or diapause time points. For most triglycerides, levels are higher at both 1-month (1m) and 6-days (6d) diapause relative to development. The bottom panel shows the fold change values of the same lipids in the African turquoise killifish compared to the red-striped killifish; levels of most triglycerides are higher in African turquoise killifish. Green boxes and text below heatmap blocks denote very long-chain fatty acids among displayed triglycerides species. (F) Lipid abundance counts for very long-chain fatty acids in triglycerides (TGs) in the African turquoise killifish during development (yellow) and diapause (blue) timepoints. Bars represent mean values in each condition error bars represent standard error of the mean (SEM). Dots represent individual replicates. *P*-values from Welch’s t-test. (G) Normalized lipid abundance counts for very long-chain fatty acids in triglycerides in the African turquoise killifish (left) and red-striped killifish (right) during matched developmental timepoints (Pre-Dia). Data represented as in F. *P*-value from Welch’s t-test. (H) Putative model for the evolution of diapause-specialized paralogs in the African turquoise killifish, in which very ancient paralogs (>473 MY) have undergone regulatory rewiring (TF binding site occurrence and chromatin-accessibility change) that generates a diapause-specialized (blue) and development-specialized (yellow) gene in the paralog pair. Some of these diapause expressed paralog genes are important for lipid metabolism, such as the processing of very long-chain triglycerides. These very long-chain triglycerides are abundant in the African turquoise killifish and accumulate in diapause, and they could be critical for long-term survival.

While some lipids and metabolites have been examined in killifish embryos and adults (*29–31*), a systematic profiling of lipids in diapause vs. development in killifish species with and without diapause has not been done. We therefore performed untargeted lipidomics on the African turquoise killifish embryos at different times: pre-diapause and diapause at different times (6 days and 1 month). As a comparison, we also performed lipidomics on embryos of another killifish species that does not undergo diapause (red-striped killifish) (Fig. 5C). The lipidome separated diapause from development in the African turquoise killifish and development in the red-striped killifish by PCA (Fig. 5D). Glycerophospholipids (e.g., phosphatidylcholines [PC]), which are membrane lipids, and triglycerides (TGs), which are storage lipids, were both changed in diapause in comparison to development stages (fig. S15A). The triglyceride changes in diapause are consistent with expression differences of genes, including specialized paralogs, encoding triglyceride metabolism enzymes and regulators (fig. S15B). Interestingly, we observed an enrichment of TGs containing very long chain fatty acids (fatty acids with chain lengths of 22) in diapause compared to development, and the majority of these very long chain fatty acids have 5 (docosapentaenoic acid, DPA) and 6 (docosahexaenoic acid, DHA) double bonds (Fig. 5, E and F). The same TGs with very long-chain fatty acids were also more abundant in African turquoise killifish embryos, even at pre-diapause, than in red-striped killifish at the equivalent state of development (Fig. 5G and fig. S15, C and D). As very long-chain fatty acids are processed by peroxisomes and subsequently by mitochondria to produce energy (*32*), they may serve as a long-term energy reserve for diapause. Other lipids, such as many ether-linked glycerophospholipids (plasmalogens), which can protect brain and hearts from oxidative stress (*33–35*), are more abundant in diapause than development and higher in the African turquoise killifish compared to the red-striped killifish (fig. S15D, Data File S8). Collectively, these data suggest that the African turquoise killifish has evolved to pack specific lipids, including long-chain fatty acids and membrane lipids with antioxidant properties, in their embryos. The rewiring of key transcription factor binding sites (e.g., FOXO1 or PPAR binding sites) at specialized paralogs (and other genes) involved in lipid metabolism could modulate lipid management for long-term protection and efficient storage and usage of specific fatty acids (Fig. 5H).

## Discussion

Our study shows for the first time that although diapause evolved recently (less than 18 MY ago), the paralogs that specialized for diapause are ancestral and shared by most vertebrates (>473 MY old). This paralog specialization in the African turquoise killifish diapause is likely achieved by recent co-opting of conserved transcription factors (such as REST, FOXOs, and PPAR) and repurposing of their regulatory landscape by mutations and selection and transposon element insertion. Our multi-omics analysis of the diapause state (transcriptomics, chromatin states, lipidomics) and comparative analysis with several fish species suggests a model for diapause evolution via very ancient paralog-specialization. After duplication in the ancestor of most vertebrates, these very ancient paralogs likely specialize in the transient response to harsh environment (e.g., transient lack of food or temperature change, or other changes), which ensures their long-term maintenance in the genome. When the ancestors of African turquoise killifish transitioned to ephemeral ponds over 18 million years (*14*), these paralogs evolved new transcription factor binding motifs driving further specialization, notably for lipid metabolism genes, for survival under extreme conditions in diapause (Fig 5H).

Elucidating the mechanisms underlying the origin of complex adaptations and phenotypes (e.g., ‘suspended animation’, novel cell types/tissues, etc.) is a central challenge of evolutionary biology (*36*). Gene duplication is the primary mechanism to generate new genes, and these act as substrate to evolve new functions. For example, ancient gene duplicates (paralogs) are specialized for expression in different tissues (*37, 38*). Ancient gene duplicates (paralogs) can also contribute to the evolution of new organs such as electric organ (*39*) and placenta (*40*). Gene duplicates have also been correlated with exceptional resistance to cancer in long-lived species (*41–43*). The specialization of paralogs might also explain how the African turquoise killifish genome can support two seemingly antagonistic complex traits – e.g., rapid life and long suspended animation. However, the mechanisms of how divergence of duplicated genes or paralogs contribute to the evolution of complex adaptations are still poorly understood. Emerging evidence, including our study, suggests that complex adaptations can arise by rewiring gene expression by unique regulatory elements (*22, 23, 44, 45*). Our results indicate that such a rewiring can be achieved using de novo regulatory elements and in some cases transposon insertion.

Cis-regulatory elements such as enhancers and promoters are known to evolve rapidly (*46–49*), and they can in turn facilitate complex adaptations with the same set of conserved genes. Transposon insertion can be even faster in promoting the rearrangement of regulatory regions (*50–57*). Rapid reshuffling of regulatory regions by mutation or transposon insertion provides a framework for the evolution of complex trait in nature. Such a mechanism could extend to the evolution of other complex traits, including regenerative capacity, which involves new enhancers in killifish (*58*), although other mechanisms may also contribute, including positive selection of specific genes (*8, 9, 26, 59–61*).

Our work also reveals specific genes and lipids that could be critical for long-term survival in suspended development. Lipids that accumulate in a state of ‘suspended animation’ (e.g., very long-chain fatty acids) could serve as key substrates for long-lasting survival (*62, 63*). Alternatively, they could also serve as new signals to affect specific aspects of the diapause state (in addition to known signals in other species, such as vitamin D or dafachronic acid in the South American killifish (*64*) and in *C. elegans,* respectively) (*65, 66*). The pathways and regulatory mechanisms we identified could also apply to other states of suspended animation and even to adult longevity. For example, transcription factors whose motifs are enriched in the diapause state (e.g., FOXOs, PPARs) are genetically required for suspended animation states, such as *C. elegans* dauer (*67–69*), and are expressed in mammalian hibernation (*70*). Furthermore, lipids and lipid metabolism genes are expressed differentially in mammalian diapause (*71*) as well as hibernation and torpor (*63, 72, 73*), and are under positive selection in exceptionally long-lived mammals (*74*). Finally, several of the transcription factors we identified (e.g., FOXOs, REST) are genetically implicated in longevity in *C. elegans* and flies (*75–80*). Our results place these previous genetic and expression findings in a natural context and reveal how selective pressure can co-opt key metabolic programs to achieve extreme phenotypes. These observations also raise the possibility that a core program of lipid metabolism genes, regulated by specific transcription factors, can be deployed to achieve metabolic remodeling and stress resistance in diverse contexts, including in adults. Our study provides a new multi-omic resource for understanding the regulation and evolution of suspended animation states (hibernation, torpor, diapause). It also opens the possibility for strategies, including lipid-based interventions, to promote long-term tissue preservation and counter age-related diseases.

## ACKNOWLEDGMENTS

We particularly thank J. Pritchard for invaluable discussion on paralogs and evolutionary analyses and for feedback on the manuscript. We thank J. Podrabsky and J. Wagner for providing A. limnaeus embryos and helpful suggestions with husbandry. We thank M. Robinson-Rechavi and J. Liu for help with positive selection analysis in ATAC-seq peaks. We thank J. Miklas, X. Zhao, K. Papsdorf, T. Ruetz, J. Chen, H. Fraser, G. Bejerano, S. Goenka, H. Chen, and all Brunet lab members for helpful discussion or comments. We thank C. Bedbrook, E. Sun, F. Boos, T. Ruetz, and X. Zhao for independent code-checking. Funding: This work was supported by the Glenn Foundation for Medical Research (A.B.), a fellowship from Stanford Center for Computational, Evolutionary, and Human Genomics (P.P.S.), and a National Science Foundation graduate fellowship (G.A.R.). Author contributions: P.P.S. designed the project with help from G.A.R. and A.B.. P.P.S. performed all the killifish experiments and computational analyses unless otherwise indicated. G.A.R. generated all ATAC-seq libraries, performed multi-genome integration of ATAC-seq and transposon analyses. K.C. and M.E. generated the lipidomics data and helped with the analysis under the supervision of M.P.S.. C-K.H. provided African turquoise killifish RNA-seq datasets pre-publication. P.P.S. wrote the manuscript with the help from G.A.R. and A.B.. All the authors provided intellectual input and commented on the manuscript.

## SUPPLEMENTARY MATERIALS

Material and Methods

References 81–131

Figs. S1 to S17

Table S1

Data Files S1 to S8

