## Supplementary material for "Evolution of diapause in the African turquoise killifish by remodeling ancient gene regulatory landscape": Materials and Methods

**This PDF file includes:**

Materials and Methods
References 81–131
Figs. S1 to S17
Table S1

**Other Supplementary Material for this manuscript includes the following:**

Data Files S1 to S8 as Excel File.

**MATERIAL AND METHODS**

All the RNA-seq and ATAC-seq data generated in this study have been deposited to NCBI-GEO (accession # GSE185817) and can be accessed at: https://www.ncbi.nlm.nih.gov/geo/query/acc.cgi?acc=GSE185817. All the lipidomics data generated in this study have been deposited to the Metabolomic Workbench (Study ID ST001898) and can be accessed at: http://dev.metabolomicsworkbench.org:22222/data/DRCCMetadata.php?Mode=Study&StudyID=ST001898&Access=NguY6247. All the code for data analysis can be accessed on GitHub at: https://github.com/param-p-singh/Diapause-multiomics

#### 1. Identification and dating of paralogs

To generate a comprehensive resource of paralogs in multiple killifish species and to date their duplication time relative to other species, we used the OrthoFinder pipeline (*15, 81*). To this end, we collected genome sequences from multiple killifish species with and without diapause from published reports and NCBI genome (*8, 9, 26*), other teleost fish, mammals, and non-vertebrate outgroups from Ensembl (version 100) (*82*) (fig. S2, B and D). Phylogenetic tree-based inference of orthologs, paralogs, and relative duplication timing of each paralog in all these species was done by OrthoFinder (fig. S2A). OrthoFinder infers orthogroups or gene families, orthologs between each species pair, the complete set of gene trees for all orthogroups, the rooted species tree, and all gene duplication events and their relative duplication time based on a phylogenetic approach (*15, 81*). For the species used in our analysis, we filtered out the paralog gene pairs with >20 partners for a gene to exclude large multigene families with inflated paralog numbers. Our results were not dependent on the paralog family size (fig. S4, D and E). Duplication node and approximate timing of the duplication (in Million Years [MY]) for each paralog pair was annotated based on known phylogenetic tree from Ensembl for species covered in Ensembl version 100 (*82*) or published reports for killifish species (*14*). To ensure that our results were not affected by the choice of species and outgroups used, we used 3 different sets of species to run the complete OrthoFinder pipeline independently: a set of 71 species, 31 species, and 13 species (fig. S2, B and D). The three pipelines resulted in very similar estimates of relative duplication time for killifish paralogs and the results were qualitatively identical (Fig. 1E and fig. S4, A to E). We used paralogs identified by OrthoFinder analysis with 71 species for our study (20,091 paralog pairs in African turquoise killifish, *Nothobranchius furzeri* and 22,955 pairs in the South American killifish, *Austrofundulus limnaeus* genomes).

In addition to OrthoFinder, we also annotated the paralog duplication timings in the African turquoise killifish directly from Ensembl version 84 using an independent approach. To identify the paralog pairs in the African turquoise killifish genome, we first identified high confidence one–to-one orthologs (bi-directional best hits) between the African turquoise killifish and each of the 5 teleost fish species (zebrafish, *Danio rerio*; medaka, *Oryzias latipes*; stickleback, *Gasterosteus aculeatus*; tetraodon, *Tetraodon nigroviridis*; and fugu, *Takifugu rubripes*) using BLASTp (E-value 1e-03) (*83*). We next identified paralogs in each of the five teleost fish for which both the genes had one-to-one orthologs in African turquoise killifish, and assigned their duplication time to the African turquoise killifish paralog. Because Ensembl did not have any killifish species, the paralogs duplicated in the killifish lineages after the divergence from medaka would be missed. Therefore, to identify such paralog pairs, we performed a protein family clustering using all the protein coding genes for multiple killifish species with and without diapause along with other teleost fish. We then annotated the duplication time for each of the potential paralogs that were not already identified using the ortholog analysis as "teleost" (if they were shared with the other teleost fish), "aplocheiloidei" (i.e. common ancestor of all killifish, if it was shared only by killifish species without diapause), and "nothobranchius" or "*Nothobranchius furzeri*" (shared by nothobranchius genus or only present in the African turquoise killifish, respectively).

To simplify the interpretation and analysis, the relative duplication nodes from each analysis were divided into 3 categories: *very ancient* (paralogs duplicated in the ancestor of jawed vertebrates at nodes Gnathostomata and earlier i.e. >473.3 MY ago), *ancient* (paralogs shared by most teleost fish species, duplicated between nodes Ovalentaria and Gnathostomata at 111-473.3 MY ago or earlier), *recent/very recent* (paralogs shared by most killifish species, duplicated between nodes Ovalentaria and *Nothobranchius furzeri* at < 111 MY ago) (*82*) (Fig 1D, figs. S2 to S4). Diapause-specialized paralog numbers (see below) in each of the three categories were compared to the genome average in that category with 10,000 bootstraps resampling of 50% paralogs genome-wide (Fig. 1E, Fig. 2E and figs. S3 and S4).

**Identification of paralogs retained from whole genome duplication (WGD).** Paralogs retained from the whole genome duplication (also called ohnologs) are known to have distinct evolutionary and genomic properties (*84*). To identify how diapause evolution is affected by WGDs, we identified ohnologs in the African turquoise killifish genome retained from the two rounds of vertebrate ancestral WGDs (which occurred around 500-550 MY ago) or the teleost (bony) fish specific third round of WGD (which occurred around ~350-400 MY ago) using multi-genome synteny comparison implemented in the OHNOLOGS database (*85, 86*). Briefly, we compared macro-synteny (gene content on chromosomes irrespective of their exact order) between African turquoise killifish and multiple outgroup genomes diverged before the respective WGD (outgroup comparison) using OHNOLOGS v2 pipeline (*86*). A similar synteny comparison was performed between the regions in the African turquoise killifish genome in a genome-wide manner (self-comparison). To identify ohnologs, we used the paralogs sets generated from co-orthology analysis in Ensembl version 84 in the African turquoise killifish as our input and identified the ones that have a significant q-score (*85*) corresponding to the relaxed criteria (outgroup q-score < 0.05 and self-comparison q-score < 0.3; http://ohnologs.curie.fr/) (*86*). This resulted in 8,810 ohnologs from vertebrate WGDs and 3,109 ohnologs from the teleost fish specific WGD (fig. S3A).

**Classifying paralogs specialized for killifish diapause.** To identify African turquoise killifish paralog pairs that show signs of specialization of the gene expression pattern for diapause, we used the normalized RNA-seq expression from Hu et al. (*6*) (see below). This dataset consists of two stages during African turquoise killifish development (heartbeat onset and diapause escaped embryos 1-day post heartbeat onset) and three time points during diapause (diapause embryos at 3 days, 6 days, and 1 month in diapause). To test robustness, we used several different criteria to identify diapause-specialized paralogs (different FDR cutoffs, and different combinations of differentially expressed genes). We first identified differentially expressed genes in all three diapause time points with respect to both development time points using DESeq2 (version 1.30.1) (*87*). A paralog gene pair was classified as having specialization of expression if one gene was significantly upregulated in one of the three diapause time points (FDR < 0.05) with respect to one of the two development stages, and the other partner gene was significantly downregulated in diapause or had a median expression in development higher than median expression in diapause. The slightly relaxed condition for development gene expression was used to maximize the number of pairs with potential specialization for downstream analysis. This resulted in 6,247 paralog pairs with expression specialization in diapause with the 71 vertebrate OrthoFinder pipeline (Data File S2).

We independently identified paralogs specialized for South American killifish diapause, using RNA-seq data of South American killifish embryos in diapause and development (4 days post diapause exit) from Wagner et al. (*9*). Paralogs with one gene significantly expressed (i.e., upregulated) in diapause compared to development (FDR < 0.05), and the other gene significantly expressed (i.e., downregulated) in development compared to diapause (FDR < 0.05) were classified as specialized paralogs (2,480 pairs).

#### 2. Killifish husbandry and embryo sample collection

The killifish and other outgroup species used in this study are listed in Table S1. All the killifish species used for data generation were housed in the Stanford Research Animal Facility II under the approved protocol (protocol #APLAC-13645). Animals were housed in automated circulating water system with pH maintained at 6-7.5 and conductivity maintained between 3500 and 4500µS/cm with a 10% system water exchange every day by reverse osmosis treated water. Adult fish were manually fed Otohime fish diet (Reed Mariculture, Otohime C1 [Ep1 for the South American killifish]) twice a day during weekdays and once a day during weekends.

Newly hatched fries for all species were kept in 0.8-liter fry tanks at a density of 4-5 fries for first two weeks and then individually housed for next two weeks. Fries were fed newly hatched brine shrimps (Brine Shrimp Direct, 454GR) twice a day during weekdays, and once a day during weekends. Animals were sexed at 4 weeks of age and transferred to 2.8-liter tanks. For African turquoise killifish and South American killifish (with diapause), adult males and females were individually housed except for breeding. Red-striped killifish, and lyretail killifish adults were kept in pairs with one male and one female animal in each tank.

For breeding, African turquoise killifish and South American killifish (with diapause) males and females were transferred to breeding tanks for a period of ~5 hours. Breeding tanks had sand trays at the bottom for the African turquoise killifish and trays with extra coarse grade glass beads (30/40 Mesh, 425-560micron size, Kramer Industries Inc. USA) for the South American killifish as per the established protocols (*6, 88-90*). After ~5 hours, sand or glass beads were filtered using a sieve to collect embryos. For the red-striped killifish and the lyretail killifish (without diapause), spawning mops constructed using green yarn were floated from the lid. The yarns were checked every day for embryos, and the embryos were carefully hand-picked.

We used young animals (1-3 months of age) for breeding and embryo collection. For each species, collected embryos were washed multiple times and live embryos were placed in Ringer's solution (Sigma-Aldrich, 96724) with 0.01% methylene blue at 26°C. Embryos were checked under a stereoscope every day and any dead embryos were removed.

**Staging of embryos.** Synchronized killifish embryos for African turquoise and South American killifish were collected within a tight (~5 hour) breeding window. Most collected embryos were at the 1-2 cell stage upon collection. We monitored embryos every day post-collection to observe the visual markers of diapause and development as previously described (*6*). Briefly, we used Kupffer's vesicle (KV), which is a transient embryonic organ present from early to middle somitogenesis as a marker to stage embryos that are about to reach diapause. KV-positive embryos reach the end of somitogenesis in 1-2 days and the loss of KV roughly coincides with the onset of heartbeat in killifish, followed by either diapause or continue development (*6, 91*). We counted the number of somites in KV-positive embryos and designated KV-positive embryos at 15-22 somites as our *"pre-diapause (Pre-Dia) stage"*. Embryo morphology for all the killifish species was similar at this stage. This mid-somitogenesis time point also coincides with the vertebrate phylotypic period (the period of the most conserved gene expression pattern during vertebrate development) with available gene expression and chromatin accessibility data from multiple other fish species (*21*).

In killifish species with diapause, young mothers have most of their embryos develop directly, whereas more mature mothers (even before middle age) have an increased frequency of embryos in diapause (*6, 92*). This feature allows us to collect *pre-diapause* embryos, even though there are no known markers, as of yet, to determine if embryos at an earlier stage are destined to diapause. Therefore, for the African turquoise and South American killifish, we collected *pre-diapause (Pre-Dia)* embryos from the very first breeding session (first clutch) from young mothers and fathers (age 4-5 weeks) with most embryos expected to skip diapause and continue developing which ensured that we get development bound embryos at *pre-diapause (Pre-Dia)* stage.

Among the first visual markers of diapause is the slowing of the rate of heartbeat after its onset (*6, 93*). Therefore, we next monitored the onset of heartbeat, and stage diapause embryos at *6 days (Dia 6d)* and *1 month of diapause (Dia 1m*) as exhibiting a continuously decreasing heartbeat rate since diapause onset (<45 beat-per-minute (BPM)) as described in Hu et al (*6*). For embryos in 1 month diapause (*Dia 1m*), we additionally made sure that there was no heartbeat by monitoring them for 5 minutes under a stereoscope to verify that they were not prematurely exiting the diapause state. For embryos in development, embryos that had an increase in heartbeat rate 1 day after heartbeat onset (>45 BPM), but before the visual pigmentation in eyes was developed (i.e. before pharyngula stage) were designated as *developing embryos (Dev)* (*6*). All the diapause and development stages stage are identical to our previous study (*6*), except *Pre-Dia* stage which is a day before the onset of heartbeat. For the South American killifish, we could only obtain one replicate for *Pre-Dia, Dev* and *Dia (1m)* stages each, due to colony loss upon facility restrictions for the COVID-19 pandemic. For killifish species without diapause (red-striped and lyretail killifish), we followed the same staging procedure described above to collect embryos at *pre-diapause (Pre-Dia) stage*. Because there is no diapause in these killifish, development embryos were taken as 1 day after the onset of heartbeat to match to the *Dev* stage in the African turquoise killifish.

**Embryo sample collection.** For each stage in each species, roughly 8-30 embryos were carefully dissected in ice-cold PBS using biological-grade tweezers (Electron Microscopy Sciences, 72700-D) to carefully remove the chorion, the enveloping layer, and the yolk without damaging the embryo body. Freshly dissected embryos were then quickly rinsed with ice-cold PBS, and all the PBS was carefully removed. Embryo bodies were then snap-frozen in liquid nitrogen and stored at -80°C. We used 8-10 snap-frozen embryos for RNA-seq and ATAC-seq and 25-30 embryos for lipidomics (see below). The details of all samples and stages used are in Data File S1.

#### 3. RNA-seq library preparation and analysis

To profile gene expression at pre-diapause stages in the African turquoise, red-striped and lyretail killifish, we constructed RNA-seq libraries (Data File S1, GSE185815, https://www.ncbi.nlm.nih.gov/geo/query/acc.cgi?acc=GSE185815). Snap frozen embryos at -80°C were thawed on ice for 1 minute and washed with 200µl ice-cold PBS. The embryos were then dissociated and homogenized with ~25 Zirconia/Silicon 0.5mm glass beads (RPI, Research Products International Corp, 9834) using FastPrep® -24 homogenizer (MB Biomedicals, 116004500) for 20 seconds, followed by centrifugation (17000g for 3 minutes). After centrifugation, 10.5µl of the supernatant was used as input to the SMART-Seq® v4 Ultra® Low Input RNA Kit (Takara, 634890) for the cDNA synthesis followed by amplification with 12 cDNA amplification cycles. Amplified cDNA was validated with Agilent 2100 Bioanalyzer using Agilent's High Sensitivity DNA Kit (Agilent, Cat. No. 5067-4626). The DNA libraries were then generated using the Nextera XT DNA Library Prep Kit (Illumina, FC-131-1096). Library quality and concentration were assessed by the Agilent 2100 Bioanalyzer and Agilent's High Sensitivity DNA Kit (Agilent Technologies, Cat. No. 5067-4626), followed by high throughput sequencing on Illumina HiSeq platform with 2 x 150bp paired end reads.

In addition, we also used available African turquoise killifish (*6*), South American killifish (*9, 64*), medaka (*21*) and zebrafish (*21, 94*) embryo RNA-seq data for our analysis (Data File S1), and processed them using the same pipeline described below. For medaka and zebrafish, we used mid-somitogenesis stages for our analysis that are expected to be the closest across vertebrates (*21*) (Data File S1).

**RNA-seq data analysis.** We first trimmed the adaptors from raw sequencing FastQ files using Trim Galore (version 0.4.5) (http://www.bioinformatics.babraham.ac.uk/projects/trim_galore/) followed by read quality assessment using FastQC (version 0.11.9, https://www.bioinformatics.babraham.ac.uk/projects/fastqc/) and MultiQC (version 1.8) (*95*). Adaptor trimmed files were aligned to the respective genomes (Table S1) using STAR (version 2.7.1a) (*96*). No reference genome is available for the red-striped killifish, so the reads from red-striped killifish RNA-seq libraries were aligned to the genome of its close relative, lyretail killifish. Identification of accurate gene expression values for paralogs can be challenging if the reads align to both the genes in the pair equally well. Therefore, we excluded all the reads that mapped to multiple locations in the genome, and only kept reads that align uniquely to a single genomic locus with samtools (version 1.5) using "*samtools view -q255*" command. Read counts were then assessed using featureCounts function in Subread package (version 2.0.1) (*97*). Raw gene expression values were then normalized using DEseq2 (version 1.30.1) (*87*). Because different

RNA-seq datasets were generated separately, we performed separate normalization for each of the individual analyses.

#### 4. ATAC-seq library preparation and analysis

To identify diapause-specific regulatory regions in the genome of African turquoise killifish and how these have evolved, we performed the Assay of Transposase Accessible Chromatin followed by high throughput sequencing (ATAC-seq) (*20, 98*) in the embryos of multiple species. ATAC-seq is an unbiased and sensitive assay of genome-wide accessible chromatin landscape that requires very low input material. We performed ATAC-seq on embryos collected from five different killifish species with and without diapause, and at different stages of development and diapause (Data File S1, GSE185816, https://www.ncbi.nlm.nih.gov/geo/query/acc.cgi?acc=GSE185816). To generate nuclei-suspension for ATAC-seq libraries, snap frozen embryo samples (~10 embryos per sample) were thawed for 1 minute and resuspended at 4°C in 200µl EZ-lysis buffer (Sigma Aldritch No. 3408). Samples were then transferred to 250µl mini-douncers (DWK (Kimble) 885300-0000) and dounced 25 times with pestle A and B respectively. After a 2 minute incubation following douncing, samples were spun at 500g for 5 minutes to precipitate nuclei, and the EZ-lysis supernatant was removed. Nuclei were then resuspended in 250µl PBS (ThermoFisher No. AM9624) and an aliquot of 5µl of nuclei was incubated with 5µl of 0.4% trypan blue stain (ThermoFisher No. 15250061) for counting the total intact nuclei counts.

Samples of ~25,000 nuclei were then suspended in a Tn5 transposition mix (65µl of tagmentation DNA buffer (Illumina No. 20034197), 63µl of nuclease-free water, and 2.5µl of tagmentation DNA enzyme I (e.g Tn5 transposase) (Illumina No. 20034197) for 20 minutes at 37°C. Following incubation, the mix was purified using the Qiagen mini-elute kit (Qiagen No. 28206) to isolate tagmented DNA. PCR amplification and subsequent qPCR monitoring was performed as described in the original ATAC-seq protocol (~14-18 cycles of PCR) (*20*). Amplified DNA from the PCR reaction was purified using the Qiagen mini-elute kit (Qiagen No. 28206), as recommended by the manufacturer. Samples were subsequently pooled and sequenced using next-generation short-read sequencing on an Illumina Nextseq 550 (Illumina No. PE-410-1001) with 75bp paired-end reads.

**ATAC-seq data analysis.** To process ATAC-seq, we first removed adaptors from FastQ files using TrimGalore (version 0.4.1) (http://www.bioinformatics.babraham.ac.uk/projects/trim_galore/), followed by read quality assessment with FastQC (version 0.11.9, https://www.bioinformatics.babraham.ac.uk/projects/fastqc/) and MultiQC (version 1.8) (*95*). Reads were then aligned to their respective reference genomes (Table S1) using BowTie2 (version 2.2.5) (*99*) with "--*very-sensitive*" option. No reference genome is available for the red-striped killifish, so the reads from red-striped killifish ATAC-seq libraries were aligned to the genome of the closest sequenced species, lyretail killifish. Duplicates were marked using Picard (version 2.22.1) (https://github.com/broadinstitute/picard). Duplicates, multimapping reads (MAPQ < 20), unmapped and mate-unmapped reads (only one read of the pair mapped), not primary alignments, and reads failing platform were then removed using SAMtools (version 1.5) (*100*). Because the Tn5 transposase binds as a dimer and inserts two adaptors separated by 9bp, all aligned read positions on + strand were shifted by +4bp, and all reads aligning to the – strand were shifted by −5bp, using alignmentSieve in deepTools (version 3.2.1) (*20, 101*). We called peaks using MACS2 (version 2.1.1.20160309) (*102, 103*) using different effective genome size for each species (e.g., genome size after removal of gaps represented by Ns).

Library quality was assessed using metrices recommended by ENCODE consortium (https://www.encodeproject.org/atac-seq/) including fragment length distribution to assess nucleosome bending patterns and enrichment of ATAC-seq peaks at transcription start sites. We observed the nucleosome bending pattern expected in ATAC-seq data in our libraries and there was a significant enrichment of ATAC-seq peaks at transcription start sites as expected (fig. S8). Therefore, except for a 1 month diapause sample for South American killifish which also had a low alignment rate of 37%, all the libraries were of high quality. Because of a single replicate and lower alignment rate of South American killifish samples (especially 1 month diapause sample), we only used South American killifish for Principal Component Analysis and internal comparison. We excluded the South American killifish samples from our genome-wide conservation analysis across species.

ATAC-seq data from medaka and zebrafish for corresponding development stages were obtained from Marlétaz et al. (*21*) (Data File S1) and processed using the same pipeline described above. We used development stage 19 and 25 in medaka and 8-somites and 48 hours post fertilization in zebrafish, which are expected to correspond to pre-diapause and development in African turquoise killifish respectively. These were used for chromatin accessibility conservation analysis presented in Figs. 3 and 4.

**Identification of diapause specific ATAC-seq peaks.** To identify ATAC-seq peaks that are specific to diapause in the African turquoise killifish genome, we performed a differential peak accessibility analysis pairwise between the two developmental conditions (pre-diapause and non-diapause) and the two diapause conditions (diapause at 6 days and 1 month time points) using DiffBind (version 2.16.2) (*104, 105*). We used both DESeq2 (*87*) and edgeR (*106*) algorithms implemented in DiffBind for differential accessibility analysis. Diapause specific peaks were then identified as the peaks that were significantly up (chromatin more open) in any of the two diapause conditions with either DESeq2 or edgeR, but do not significantly change (up or down) between the two development conditions with both DESeq2 and edgeR. This led to 6,490 chromatin peaks genome-wide in African turquoise killifish that are significantly up in diapause but do not change during development (fig. S7A, Data File S3). Peaks were assigned to their nearest genes using ChIPseeker (version 1.28.3) (*107*), to identify 1,880 diapause specific peaks at specialized paralogs (Data File S3). Peak annotation with the genomic properties was also performed using ChIPseeker (fig. S7B). These peaks at specialized paralogs were used for motif enrichment and peak conservation analyses presented in Figs. 3 and 4.

#### 5. Multiple whole-genome alignment

To integrate ATAC-seq and RNA-seq datasets across species, we performed a 5-way multiple whole-genome alignment with African turquoise killifish (*Nfur: Nothobranchius furzeri*), lyretail killifish (*Aaus: Aphyosemion australe*), South American killifish (*Alim: Austrofundulus limnaeus*), medaka (*Olat: Oryzias latipes*) and zebrafish (*Drer: Danio rerio*) (Table S1), using African turquoise killifish as the reference genome. For red-striped killifish (*Aphyosemion striatum*), genome of the closest sequenced species lyretail killifish was used for integrative analysis. For genomes with chromosome level assemblies, we discarded scaffolds not placed on chromosomes. First, we performed pairwise alignments between African turquoise killifish and each of the four other fish genomes using LASTZ (*108*) (parameters: --gap=400,30 --gappedthresh=3000 --

ydrop=6400 --inner=2000 --hspthresh=1500 --masking=50 --notransition --step=20 --scores=HoxD55.q). Subsequent chaining and netting were performed using the suite of UCSC genome browser utilities (*109*). The percentage of aligned African turquoise genome to each of the other fish species decreased based on the distance to the last common ancestor as expected (*110*) with 61.1%, 47.8%, 23.2%, 20.1% of the African turquoise killifish genome aligning to the lyretail killifish, South American killifish, medaka and zebrafish genomes respectively in a pair-wise manner.

These pairwise alignments were then merged using the multi-alignment tool Multic/TBA (*111*), using the command *<tba + E=Nfur ((((Nfur Aaus) Alim) Olat) Drer ./pairwise_dir/>* to obtain a single, 5-way, multiple whole-genome alignment using the African turquoise killifish genome as the reference (specified by *E=Nfur*). The resulting multiple-whole genome alignment covered ~75.3% of the African turquoise killifish genome. Coverage of each of the aligned fish genome in the multi-alignment also diminished as time to the last common ancestor increased with 62.7%, 85.9%, 14.2%, and 23.7% of the genome being covered for lyretail killifish, South American killifish, medaka, and zebrafish genomes respectively.

To assess the quality of our genome alignment, we compared the length of aligned sequence blocks in multi-genome alignment with that of teleost fish 8-way multi-genome alignments available from the UCSC genome browser and generated using a similar approach (*112*) (https://hgdownload.soe.ucsc.edu/goldenPath/danRer7/multiz8way/). We found that the aligned block lengths in both our and 8-way multi-genome alignment from UCSC were comparable. Most of the aligned blocks were either 10-99bp long (53% our vs 38.7% UCSC-fish) or 100-999bp long (33% our vs 26.5% 8-way alignment from UCSC) in both the alignments. Importantly, a vast majority of our ATAC-seq peaks (98.35% of chromosomal peaks) fall in the regions that are covered in our multi-genome alignment.

#### 6. Integrating ATAC-seq peaks across species

The 5-way multiple whole-genome alignment was used to compare ATAC-seq data across species. Bed files for each ATAC-seq library were cross-referenced to the alignment and the coordinates of ATAC-seq peaks for all species were converted to African turquoise killifish genome coordinates. During this process, peaks were tagged as “conserved” at three levels of stringency: relaxed (any base pair overlap between peaks), strict (25% of the African turquoise killifish peak must be covered by aligned peak region in other species), and very strict (50% of the African turquoise killifish peak must be covered by the aligned peak in other species). The differences in peak conservation between relaxed and strict definitions was minimal (fig. S11). Thus, subsequent analysis was performed with the relaxed peak set. During coordinated conversion, some peaks for species other than African turquoise killifish became split between two or more locations in the African turquoise killifish genome. We also included these split location peaks in our analyses. However, split location peaks represent only a minority of recovered peaks (5.2%) and are unlikely to influence our analyses.

With this finalized peak set, we then categorized each peak in African turquoise killifish and its underlying sequence into one of three conservation categories: *ancient/very ancient, recent, and very recent*. 1) Peaks considered *very recent* had only a peak in the African turquoise killifish (likely originated after divergence from killifish species without diapause at < 17.79 MY) (*14*)). 2) Peaks considered *recent* had overlapping peaks in African turquoise killifish and at least one other African killifish (i.e. lyretail killifish or red-striped killifish), but not in outgroups (medaka and zebrafish; likely originated between 17.79- 93.2 MY) (*14, 82*). 3) Peaks considered *ancient/very ancient* had overlapping peaks in African turquoise killifish, at least one other African killifish (i.e. lyretail killifish or red-striped killifish), and at least one outgroup fish (i.e. medaka or zebrafish; likely originated > 93.2 MY) (*82*) (Data File S3). To avoid confounding peaks within our *very recent* category, peaks present in the African turquoise killifish, absent in other African killifish, yet present in either zebrafish or medaka were subsequently added to the *ancient/very ancient* category despite being just outside of the above parameters. The same criteria were used to define sequence conservation. However, instead of requiring accessible-chromatin overlap, sequences were evaluated for having an aligned orthologous region in each species.

To visualize these peaks across species, we used the Integrative Genomics Viewer (IGV) (*113*). For each species, RPKM-normalized read counts were used either directly (paralog displays) or summed across replicates and across developmental/diapause stages (for single displays) to create single coverage tracks for fish without diapause and two tracks (one diapause, one development) for fish with diapause. Tracks from each species were then anchored to each other via a single conserved base in the multiple-whole-genome-alignment and extended to the exact same window size in all species. The anchor point for each peak region was chosen based on its proximity to the summit of the peak in the African turquoise killifish. Track height for each species was set automatically by IGV using either the height of the peak of interest, or, in species without a conserved peak, to the height of the tallest peak within 40kb of the anchoring base pair. These visualizations illustrate the conservation and specialization states described above.

These analyses revealed that for the majority of peaks, the genome sequences under chromatin accessible peaks are 'alignable' (i.e. conserved enough to establish orthology at the genome-wide level), but chromatin accessibility at those regions evolved very recently and exclusively in the African turquoise killifish. This pattern was consistent for genome-wide chromatin (fig. S10A) and specifically among diapause-accessible peaks at very ancient paralogs previously identified in Fig. 1. The sequence conservation is also strongest at coding sequence (exons) and decays as expected across promoters, UTRs, introns, and intergenic regions (fig. S10B).

#### 7. Principal Component Analysis (PCA) on ATAC-seq

To explore the global relationships between killifish ATAC-seq samples, we performed principal component analysis (PCA) using ATAC-seq peak intensities (normalized aligned read counts for each peak). To this end, we first generated peak intensity matrices for each of the following comparisons: 1) for the African turquoise killifish diapause and development samples (Fig. 3B, left panel); 2) for all killifish species (African turquoise killifish, South American killifish, lyretail killifish and red-striped killifish, Fig. 3B, middle panel); 3) killifish with diapause (African turquoise killifish and South American killifish, Fig. 3B, right panel). For each comparison, the peak matrix contained VST-normalized peaks intensities for all consensus peaks detected in all the samples in that comparison. Cross-species comparison only included the peak(s) conserved in all samples. The total peaks used for PCA were 60,359 for the African turquoise killifish, 1,293 for all killifish, and 3,721 for killifish with diapause. PCA plots were done using autoplot command in ggfortify (version 0.4.11) package in R (version 3.6.2).

#### 8. Motif enrichment and conservation analysis

HOMER (version 4.10), was used for transcription factor binding motif enrichment analysis (*24*), using the ATAC-seq peaks that are significantly up in diapause and were in proximity to the diapause specific paralogs for the African turquoise killifish and their orthologous conserved peaks in other species (see below). Genomes of all the species were added to HOMER using "*loadGenome.pl"* utility with the genome fasta and GFF files as input (Table S1). We then used the genomic coordinates from thebed file for the diapause specific ATAC-seq peaks at paralogs as input to "*findMotifsGenome.pl"* and specified vertebrate motifs by "*-mset vertebrates"*. Known motifs in "*knownResults.txt*" generated by the HOMER output was used for all the analyses. To remove redundancy in motifs, we performed a motif clustering using tomtom utility in the MEME suit (version 5.3.0) (*114, 115*) using the following parameters: *-thresh 1e-5 -evalue -min-overlap 6*. The resulting clusters were manually curated, and motifs were assigned to the genes coding for the transcription factors.

**Identification of conserved transcription factor binding motifs across species.** To assess the evolution and conservation of African turquoise killifish diapause-specific transcription factor binding motifs at specialized paralogs in other species, we extracted sequences of these motifs from African turquoise killifish and the corresponding aligned sequences in other species from our 5-way multiple whole-genome alignment. We observed that a vast majority of transcription factor binding motifs that are enriched in ATAC-seq peaks up in diapause at specialized paralogs in the African turquoise killifish are aligned in other species with motif-like sequences (i.e. sequences similar to the canonical motifs). To assess if these motif-like sequences are likely to be bound by their respective transcription factors, we subjected motif or motif-like sequences to a binding likelihood calculation identical to that used by HOMER (*24*). We then determined if motif-like sequences in species other than African turquoise killifish met the log odds detection threshold (defined as the $\log(X_1/0.25) + \log(X_2/0.25) + \ldots \log(X_n/0.25)$ where X is the probability of a given base being present at a given location in a given motif) computed by HOMER (*24*) during motif enrichment, which is used to determine likelihood of transcription factor bound vs. unbound sites. We also excluded motif sites in peaks where an identical motif was found near the aligned region in another species. This allowed us to detect cases where the sequence directly aligned to a motif is not conserved, but the motif is present nearby and possibly providing similar regulatory potential.

These analyses revealed that a very low number of motif-like sequences in other species are expected to bind the transcription factor at that position and can be considered as conserved transcription factor binding sites across species (4.77% on average). Thus, the vast majority of these motif-like sequences were likely used as ‘substrates’ during evolution for mutation and selection of canonical motif sequences for binding of transcription factors (Fig. 4, E and F, and fig. S12).

**Differences between singletons and paralogs.** To assess the differences between singleton genes (genes with no known paralog) and paralogs, we first identified genes that were unambiguously identified as paralogs or singletons in all the four paralog identification pipelines (see above). This led to 4,009 singleton genes and 10,069 genes with at least one paralog in the African turquoise killifish genome. We observed that singletons and paralogs both were equally likely to be upregulated in diapause using RNA-seq data compared to their genomic average (fig. S16A). However, there were differences in the regulatory motif landscape at specialized paralogs and singleton genes. Although several motifs such as REST, FOXO3 and PPARG etc. were enriched at both singletons and paralogs, the majority of the motifs were only enriched either at paralogs or at singletons (fig. S16B). For example, some transcription factor binding motifs such as TEAD2, FOXA2, JUNB etc. were only enriched at paralogs while others such as MYC, RUNX2, ETS1 etc. were only enriched at singletons (fig. S16B). This observation suggests that in the African turquoise killifish, the diapause regulatory landscape is remodeled genome-wide, but there are differences in the regulatory repertoire of singletons and paralogs.

**Conservations of motifs in alignment-independent manner.** To test that our results are not affected by multiple whole-genome alignment and our criteria to establish homology in non-coding regions, we also used another orthogonal approach to identify if diapause specific motifs at paralogs in African turquoise killifish were conserved in other species. To this end, we compared diapause specific ATAC-seq peaks at diapause-specialized paralogs in the African turquoise killifish to the corresponding peaks at their ortholog genes in other species in a genome-alignment independent manner. To establish homology independent of genome alignment, we only focused on ATAC-seq peaks at promoters of the ortholog genes (identified using protein sequences) in African turquoise killifish with diapause and in lyretail and red-striped killifish without diapause in a pairwise manner, followed by motif enrichment analysis in the respective genome. Although this excludes many potential distal enhancer elements, the results were similar to those using our multi-genome alignment (see above), and corroborate that diapause-specific transcription factor binding motifs are only present in the African turquoise killifish genome (fig. S17).

#### 9. Transposable element identification and analysis

To evaluate the contribution of Transposable Elements (TEs) for the evolution of diapause, we first developed a comprehensive map of abundance and genomic location of all TEs in the aforementioned teleost fish species used to construct the genome multi-alignment. We employed RepeatMasker (version 4.0) (*116*) to identify repetitive sequences using the *Teleosti* suite of know repeat elements <*Repeatmasker -a -s -species 'Teleostei' Input.fa*> and < *processRepeats -xsmall RMoutput.fa.gz*> allowing for a standardized repetitive element set across species. We detected similar abundances of TE classes and families as previously reported by various sources (*117*). We then identified overlap between all ATAC-seq peak coordinates and TE coordinates in African turquoise killifish. We evaluated TE enrichment at ATAC-seq peaks up specifically in diapause as compared to: 1) 'Genome': TE representation genome-wide (Fig 4H, upper panel), 2) 'Chromatin': TE representation within all ATAC-seq peaks (Fig 4H, middle panel), 3) 'Control loci': size-matched regions 10kb downstream of  ATAC-seq up specifically in diapause (Fig 4H, lower panel), using a binomial test (Mutational Patterns Package version 3.2.0) (*118*). Several TE families showed enrichment specific to differentially accessible chromatin sites specific to diapause, such as Crypton-A (DNA), Zisupton (DNA), RTE-X (LINE), and tRNA-Mermaid (SINE) (Fig 4H).

We then evaluated the overlap between these TE instances and enriched transcription factor binding motifs detected in our analysis above. These chromatin-accessible TE-embedded motifs were also evaluated for conservation across species by assessing whether 1) the TE is present at aligned location in the genome alignment and contains the transcription factor binding motif sequence, 2) the TE is present at the aligned location in other species, but lacks the transcription factor binding motif sequence, 3) the TE is absent at the aligned location, but a transcription factor binding motif still exist at this location in the alignment, or 4) both the TE and transcription factor binding motif binding site are absent at the aligned location in the other species. This analysis revealed that a majority of TE sites are exclusive to African turquoise killifish, as can be expected given the rapid rate at which the TE landscape changes (*119-121*) and given the recent TE expansion in the African turquoise killifish genome (*26*) (Fig. 4I and fig. S14).

#### 10. Positive selection analysis

**Positive selection of regulatory regions.** To evaluate whether diapause-accessible chromatin peaks show any signature of positive selection, we used a recently developed method to detect positive selection at transcription factor binding sites and accessible chromatin (*25, 122*). We scanned for signature of positive selection at the genomic DNA underlying ATAC-seq peaks with respect to: 1) ancestor of all killifish species in our analysis ('killifish ancestor'); and 2) ancestor of killifish and medaka ('pre-medaka ancestor') (fig. S13A). We first inferred ancestral sequences for these two nodes within the teleost lineage using the PAML package (version 4.8) (*123*). Alignment blocks from our 5-way fish multiple whole-genome alignment that were at least 50bp long and covered at least 50% of the ATAC-seq peaks were used for the ancestor generation and positive selection analysis. We excluded ATAC-seq peaks that were in exons to focus on regulatory elements. The ancestral sequences and the African turquoise killifish sequences were used to generate Support Vector Machine (SVM) kmer weights and positive selection was detected using hightail test as recommended (*25, 122*) (https://github.com/ljljolinq1010/A-robust-method-for-detecting-positive-selection-on-regulatory-sequences/). The Benjamini-Hochberg procedure was used for multiple hypothesis correction, and ATAC-seq peaks with FDR $< 0.1$ for either pre-killifish or pre-medaka ancestors were considered to be under positive selection (see Data File S3).

In total, we detected 3,836 and 3,928 ATAC-seq peaks with signature of positive selection using the 'killifish ancestor' and 'pre-medaka ancestor' inferred sequences respectively, with both having a strong overlap of 3,370 (76.7%) (fig. S13B). We used the union of the two groups for the downstream analysis. A total of 172 diapause-specific ATAC-seq peaks at specialized paralogs showed signature of positive selection (Fig. 4G, Data File S3). These were enriched for several of the transcription factor binding motifs detected in our previous analysis, including REST, FOXO3 and PPARs (Fig. 4G and fig. S13C). The functional enrichment of ATAC-seq peaks also included several functions related to lipid metabolism (Data File S6). These results suggest that at least a portion of genomic loci underlying diapause-specific ATAC-seq peaks may have evolved due to positive selective pressure at these loci.

**Positive selection on protein-coding gene sequences.** The protein-coding genes under positive selection in African turquoise killifish were identified using phylogenetic analysis involving 19 fish species with and without diapause as described in Wagner et al. (*9*). Briefly, protein sequences were clustered using Proteinortho (version 5.11) (*124*), followed by filtering of clusters and alignment of coding sequences of the filtered clusters using PRANK v.140603 (*125*). The resulting codon aware alignments were filtered with GUIDANCE v2.0 (*126*) to remove low quality regions. Proteins and individual amino acids under positive selection were then identified in the ancestor of African killifish with diapause (in the branch leading to the African killifish genus nothobranchius after separation from the African killifish without diapause *A. striatum*) using the branch-site model in CODEML implemented in the Phylogenetic Analysis by Maximum Likelihood package (PAML) (*123*). This ancestral branch co-insides with the time period at which evolution of diapause likely occurred in African turquoise killifish (~18 MY ago). Proteins with a *P*-value of the branch-site test less than 0.05 (without any FDR correction to maximize the number of proteins with potential signals of selection) were then filtered. This led to a list of 213 protein-coding genes under positive selection in the ancestor of killifish species with diapause after divergence from killifish species without diapause and outgroup fish species (see Data File S4).

#### 11. Functional enrichment analysis

To perform functional enrichment analysis for diapause specific African turquoise killifish ATAC-seq peaks or upregulated genes in diapause, we used Gene Ontology (GO) analysis using GOstats package (version 2.56.0) (*127*). GO terms from human and zebrafish were assigned to their killifish orthologs (best hit protein with BLASTp E-value >1e-3). For GO enrichment analysis using diapause specific ATAC-seq peaks, we used the non-redundant list of genes closest to the peaks (Data File S3) with all protein coding genes as background and performed a hypergeometric test implemented in GOstats. Similarly, for RNA-seq, we used genes upregulated in diapause (Data File S2). GO terms enriched in both diapause RNA-seq and ATAC-seq are in Data File S5, which included many GO terms related to lipid metabolism (Fig. 5, A and B, and Data File S5). We also performed GO enrichment analysis for the subset of ATAC-seq peaks that show signatures of positive selection (see above, Data File S3), and observed that several lipid metabolism related functions are enriched in the genes next to the chromatin accessibility regions that have evolved under positive selection (Data File S6).

To identify the upstream regulators of genes upregulated during diapause in the African turquoise killifish, we used Ingenuity Pathway Analysis (IPA) upstream regulator analysis (QIAGEN, March 2021 release) (Data File S7).

#### 12. Untargeted lipidomics by LC-MS and analysis

Lipidomics experiments were performed using ~30 embryos for each stage of diapause and development from African turquoise and red-striped killifish (3-4 replicates for each stage) (Fig. 5C) as previously described (*128, 129*).

Lipids were extracted in a randomized order via biphasic separation with cold methyl tert-butyl ether (MTBE), methanol and water. Briefly, 260µl of methanol and 40µl of water were added to the embryos and vortexed for 20 seconds. A lipid internal standard mixture was spiked in each sample (EquiSPLASH LIPIDOMIX, Avanti Polar Lipids (cat #: 330731), and d17-Oleic acid, Cayman chemicals (cat #: 9000432) to control for extraction efficiency, evaluate LC-MS performance and estimate concentrations of individual lipids. Samples were diluted with 1,000µl of MTBE, vortexed for 10 seconds, sonicated for 30 seconds three times in a water bath, and incubated under agitation for 30 minutes at 4°C. After addition of 250µl of water, the samples were vortexed for 1 minute and centrifuged at 14,000g for 5 minutes at 20°C. The upper phase containing the lipids was collected and dried down under nitrogen. The dry extracts were reconstituted with 150µl of 9:1 methanol:toluene.

Lipid extracts were analyzed in a randomized order using an Ultimate 3000 RSLC system coupled with a Q Exactive mass spectrometer (Thermo Fisher Scientific) as previously described (*129*). Each sample was run twice in positive and negative ionization modes and lipids were separated using an Accucore C30 column 2.1x150mm, 2.6µm (Thermo Fisher Scientific) and mobile phase solvents consisted in 10mM ammonium acetate and 0.1% formic acid in 60/40 acetonitrile/water (A) and 10mM ammonium acetate and 0.1% formic acid in 90/10 isopropanol/acetonitrile (B). The gradient profile used was 30% B for 3min, 30–43% B over 5min, 43–50% B over 1min, 55–90% B over 9min, 90-99% B over 9min and 99% B for 5min. Lipids were eluted from the column at 0.2ml/min, the oven temperature was set at 30°C, and the injection volume was 5µl. Autosampler temperature was set at 15°C to prevent lipid aggregation.

LC-MS peak extraction, alignment, quantification, and annotation was performed using LipidSearch software version 4.2.21 (Thermo Fisher Scientific). Lipids were identified by matching the precursor ion mass to a database and the experimental MS/MS spectra to a spectral library containing theoretical fragmentation spectra. The following lipid ions were used for quantification: [M+H]+ for ceramides (Cer), (lysophosphatidylcholine) LPC, phosphatidylcholine (PC), monoglycerides (MG) and sphingomyelins (SM); [M-H]- for phosphatidylethanolamines

(PE), phosphatidylinositols (PI), phosphatidylserines (PS), phosphatidylglycerols (PG) and lysophosphatidylethanolamine (LPE); and [M+NH4]+ for cholesterol ester (ChE), diglycerides (DG) and triglycerides (TG). To reduce the risk of misidentification, MS/MS spectra from lipids of interest were validated as follows: 1) both positive and negative mode MS/MS spectra match the expected fragments, 2) the main lipid adduct forms detected in positive and negative modes agree with the lipid class identified, 3) the retention time is compatible with the lipid class identified and 4) the peak shape is acceptable. The fragmentation pattern of each lipid class was experimentally validated using lipid internal standards.

Single-point internal standard calibrations were used to estimate absolute concentrations for 431 unique lipids belonging to 14 classes using one internal standard for each lipid class. Importantly, we ensured linearity within the range of detected endogenous lipids using serial dilutions of internal standards spanning 4 orders of magnitude. Median normalization (excluding TG and DG) was employed on lipid molar concentrations to correct for differential quantity of starting material. Importantly, we verified that median lipid signal (excluding TG and DG) correlated well (Pearson's correlation coefficient = 0.48, $P = 0.005$) with the total protein content in each sample as measured by the BCA Protein Assay Kit (Pierce, cat# 23225) from precipitated proteins following the biphasic separation, suggesting good sample quality. One development (diapause escape) sample had an unexpectedly low protein concentration and thus was discarded. Lipid molar concentrations for a given class were calculated by summing individual lipid species molar concentrations belonging to that class. Fatty acid composition analysis was performed in each lipid class. Fatty acid composition was calculated by taking the ratio of the sum molar concentration of a given fatty acid over the sum molar concentration across fatty acids found in the lipids of the class. Subsequently, saturated fatty acids (SFA), mono-unsaturated fatty acids (MUFA) and poly-unsaturated fatty acids (PUFA) were grouped together for comparative analysis.

Principal Component Analysis (PCA) was performed using all the lipids identified for: 1) African turquoise killifish diapause and development samples (Fig. 5D left panel); and 2) African turquoise and red-striped killifish pre-diapause samples (Fig. 5D right panel). The total of 431 filtered and normalized lipid intensities were used for PCA (see below), which were also plotted using autoplot function in ggfortify package (version 0.4.11) in R (version 4.0.5).

Discriminant analysis was performed using a Welch's t-test that does not assume equal population variances for each lipid among the two diapause (6 days and 1 month) and the two development conditions (pre-diapause and diapause escape). Lipids that were significantly different (Welch's t-test, $P < 0.05$ after multiple hypothesis correction using Benjamini-Hochberg method) between diapause and development but did not significantly change between the two development conditions were categorized as diapause specific lipids. These constitute lipids that go up or down when embryos enter diapause but do not change among the two development time points. This led to 350 diapause specific lipid changes, 80 of which were triglycerides, including very long-chain fatty acid triglycerides (Fig. 5, E to G, fig. S15, and Data File S8).

### SUPPLEMENTARY FIGURES

#### Figure S1

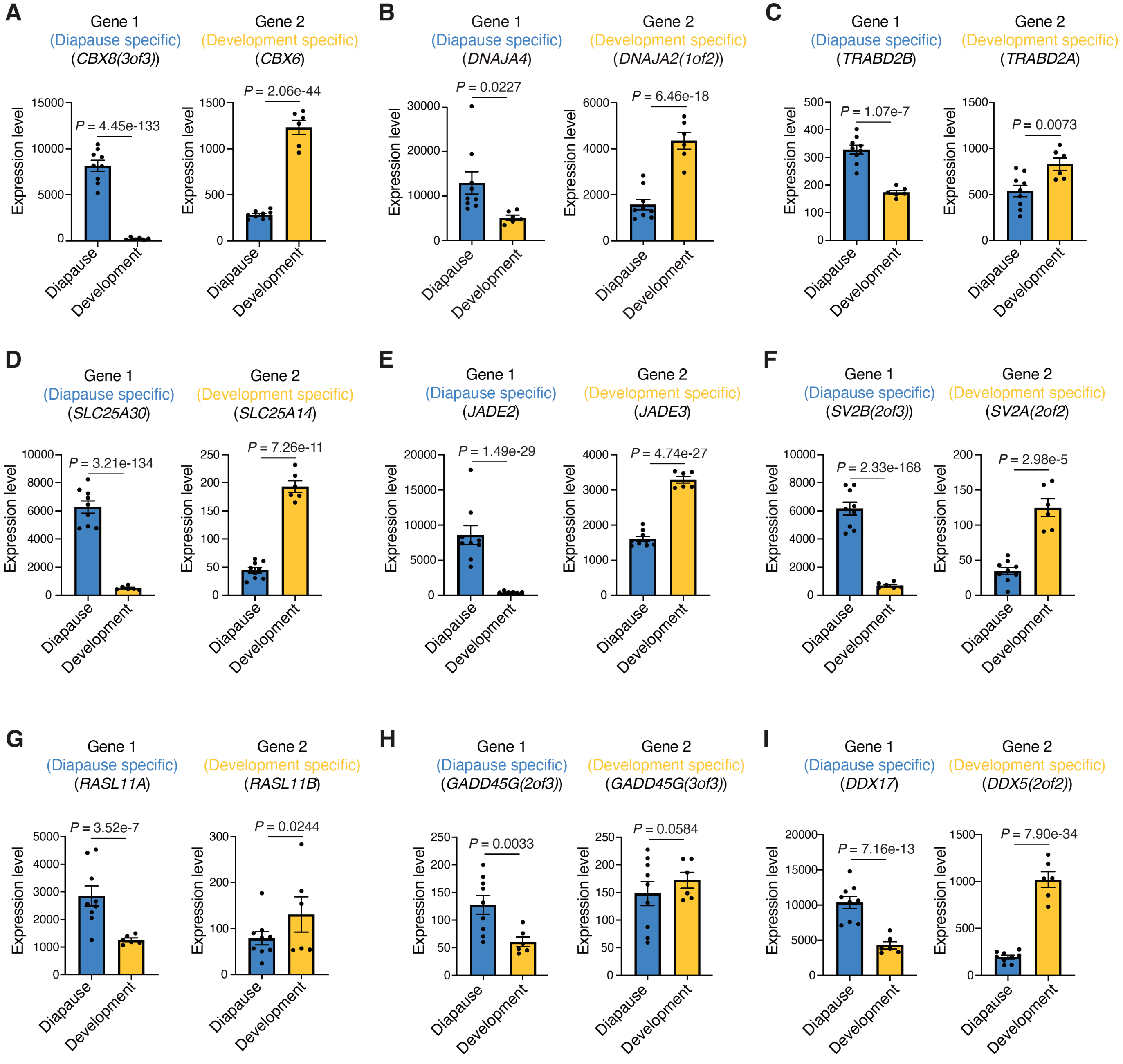

**Figure S1. Additional examples of diapause-specialized paralogs in the African turquoise killifish** (A-I) Examples of paralog gene pairs, with specialized expression of gene 1 in diapause (blue) and gene 2 in development (yellow) in African turquoise killifish (*Nothobranchius furzeri*). Bars represent mean expression level (normalized DESeq2 count) across replicates in diapause or development state. Dots show normalized DESeq2 counts in each replicate. Error bar is standard error of mean (SEM). Corrected *P*-values (median from pairwise comparisons) from DESeq2 Wald test.

Figure S2

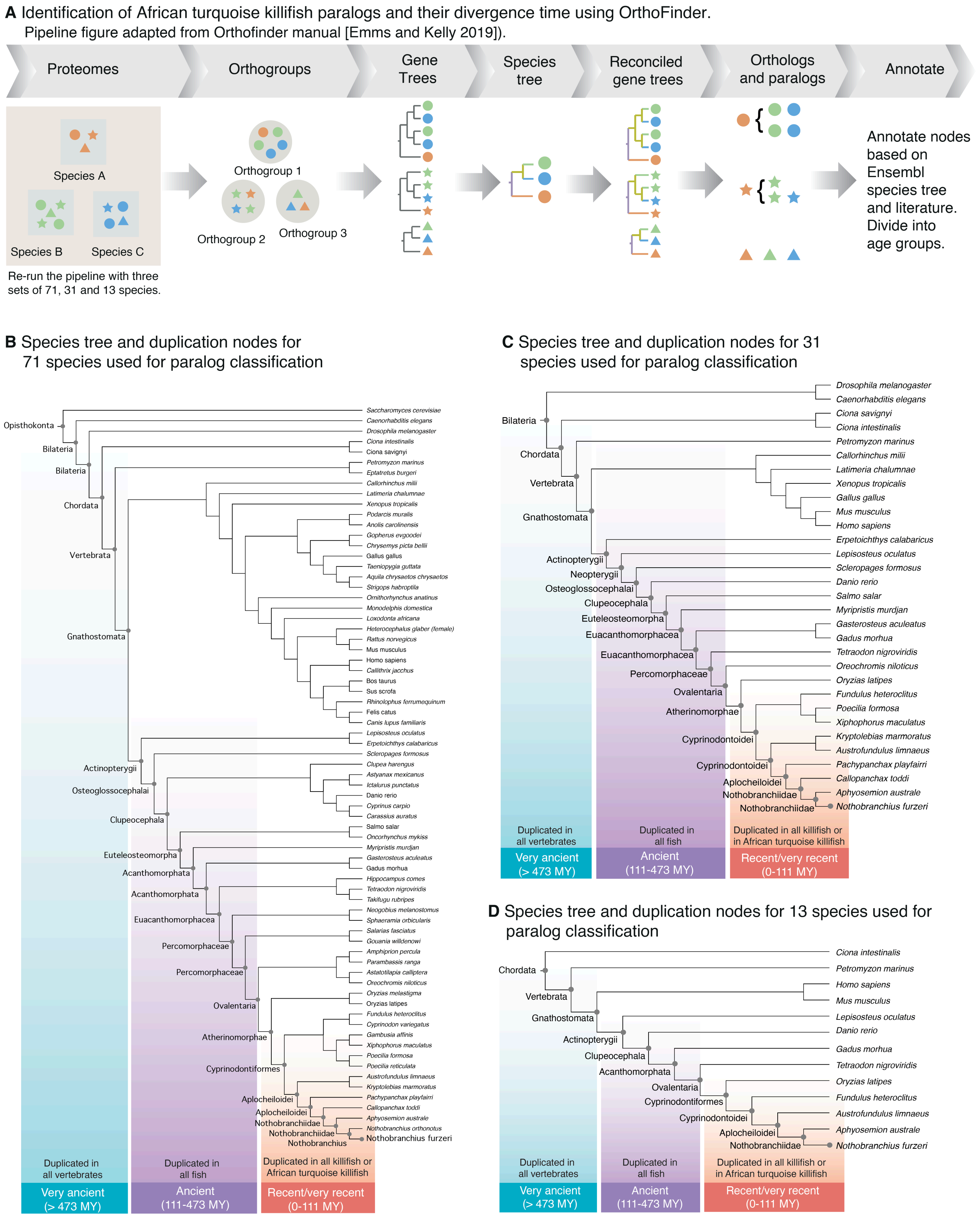
A Identification of African turquoise killifish paralogs and their divergence time using OrthoFinder.
Pipeline figure adapted from Orthofinder manual [Emms and Kelly 2019]).
Proteomes
Orthogroups
Gene Trees
Species tree
Reconciled gene trees
Orthologs and paralogs
Annotate
Species A
Species B
Species C
Re-run the pipeline with three sets of 71, 31 and 13 species.
Orthogroup 1
Orthogroup 2
Orthogroup 3
Annotate nodes based on Ensembl species tree and literature. Divide into age groups.
B Species tree and duplication nodes for 71 species used for paralog classification
Opisthokonta
Bilateria
Bilateria
Chordata
Vertebrata
Gnathostomata
Actinopterygii
Osteoglossocephalai
Clupeocephala
Euteleosteomorpha
Acanthomorphata
Acanthomorphata
Euacanthomorphacea
Percomorphaceae
Percomorphaceae
Ovalentaria
Atherinomorphae
Cyprinodontiformes
Aplocheiloidei
Aplocheiloidei
Nothobranchiidae
Nothobranchiidae
Nothobranchius
Saccharomyces cerevisiae
Caenorhabditis elegans
Drosophila melanogaster
Ciona intestinalis
Ciona savignyi
Petromyzon marinus
Eptatretus burgeri
Callorhinchus milii
Latimeria chalumnae
Xenopus tropicalis
Podarcis muralis
Anolis carolinensis
Gopherus evgoodei
Chrysemys picta bellii
Gallus gallus
Taeniopygia guttata
Aquila chrysaetos chrysaetos
Strigops habroptila
Ornithorhynchus anatinus
Monodelphis domestica
Loxodonta africana
Heterocephalus glaber (female)
Rattus norvegicus
Mus musculus
Homo sapiens
Callithrix jacchus
Bos taurus
Sus scrofa
Rhinolophus ferrumequinum
Felis catus
Canis lupus familiaris
Lepisosteus oculatus
Erpetoichthys calabaricus
Scleropages formosus
Clupea harengus
Astyanax mexicanus
Ictalurus punctatus
Danio rerio
Cyprinus carpio
Carassius auratus
Salmo salar
Oncorhynchus mykiss
Myripristis murdjan
Gasterosteus aculeatus
Gadus morhua
Hippocampus comes
Tetraodon nigroviridis
Takifugu rubripes
Neogobius melanostomus
Sphaeramia orbicularis
Salarias fasciatus
Gouania willdenowi
Amphiprion percula
Parambassis ranga
Astatotilapia calliptera
Oreochromis niloticus
Oryzias melastigma
Oryzias latipes
Fundulus heteroclitus
Cyprinodon variegatus
Gambusia affinis
Xiphophorus maculatus
Poecilia formosa
Poecilia reticulata
Austrofundulus limnaeus
Kryptolebias marmoratus
Pachypanchax playfairii
Callopanchax toddi
Aphyosemion australe
Nothobranchius orthonotus
Nothobranchius furzeri
Duplicated in all vertebrates
Duplicated in all fish
Duplicated in all killifish or African turquoise killifish
Very ancient (> 473 MY)
Ancient (111-473 MY)
Recent/very recent (0-111 MY)
C Species tree and duplication nodes for 31 species used for paralog classification
Bilateria
Chordata
Vertebrata
Gnathostomata
Actinopterygii
Neopterygii
Osteoglossocephalai
Clupeocephala
Euteleosteomorpha
Euacanthomorphacea
Euacanthomorphacea
Percomorphaceae
Ovalentaria
Atherinomorphae
Cyprinodontoidei
Cyprinodontoidei
Aplocheiloidei
Nothobranchiidae
Nothobranchiidae
Drosophila melanogaster
Caenorhabditis elegans
Ciona savignyi
Ciona intestinalis
Petromyzon marinus
Callorhinchus milii
Latimeria chalumnae
Xenopus tropicalis
Gallus gallus
Mus musculus
Homo sapiens
Erpetoichthys calabaricus
Lepisosteus oculatus
Scleropages formosus
Danio rerio
Salmo salar
Myripristis murdjan
Gasterosteus aculeatus
Gadus morhua
Tetraodon nigroviridis
Oreochromis niloticus
Oryzias latipes
Fundulus heteroclitus
Poecilia formosa
Xiphophorus maculatus
Kryptolebias marmoratus
Austrofundulus limnaeus
Pachypanchax playfairii
Callopanchax toddi
Aphyosemion australe
Nothobranchius furzeri
Duplicated in all vertebrates
Duplicated in all fish
Duplicated in all killifish or in African turquoise killifish
Very ancient (> 473 MY)
Ancient (111-473 MY)
Recent/very recent (0-111 MY)
D Species tree and duplication nodes for 13 species used for paralog classification
Chordata
Vertebrata
Gnathostomata
Actinopterygii
Clupeocephala
Acanthomorphata
Ovalentaria
Cyprinodontiformes
Cyprinodontoidei
Aplocheiloidei
Nothobranchiidae
Ciona intestinalis
Petromyzon marinus
Homo sapiens
Mus musculus
Lepisosteus oculatus
Danio rerio
Gadus morhua
Tetraodon nigroviridis
Oryzias latipes
Fundulus heteroclitus
Austrofundulus limnaeus
Aphyosemion australe
Nothobranchius furzeri
Duplicated in all vertebrates
Duplicated in all fish
Duplicated in all killifish or in African turquoise killifish
Very ancient (> 473 MY)
Ancient (111-473 MY)
Recent/very recent (0-111 MY)

**Figure S2. Identification of paralogs and their duplication timing.** (A) Schematic of pipeline used to identify and date the time of duplication of paralogs in the African turquoise killifish as adapted from the OrthoFinder manual (*34*). The proteomes of included species are grouped by protein sequence similarity and converted to gene trees. Trees for each orthogroup are then compared and reconciled against the established phylogenetic tree and used to build inferred ortholog-paralog groupings. These groups are used to identify paralogs and estimate the relative timing of gene duplication. (B) Complete dendrogram of included species for binning paralog duplication time into 3 categories based on OrthoFinder pipeline with 71 species. Divergence time estimates are from species tree in Ensembl. Binned categories are: Genes that were duplicated in the common ancestor of all vertebrates or earlier (very ancient, > 473 million years [MY]), genes that were duplicated in the common ancestor of all fish (ancient, 111-473 MY), and genes that were duplicated in the common ancestor of all killifish or African turquoise killifish exclusively (recent/very recent, 0-111 MY). (C) Complete dendrogram of included species for binning paralog duplication time into 3 categories based on OrthoFinder pipeline with 31 species. Divergence time estimates are from species tree in Ensembl. (D) Complete dendrogram of included species for binning paralog duplication time into 3 categories based on OrthoFinder pipeline with 13 species. Divergence time estimates are from species tree in Ensembl.

Figure S3

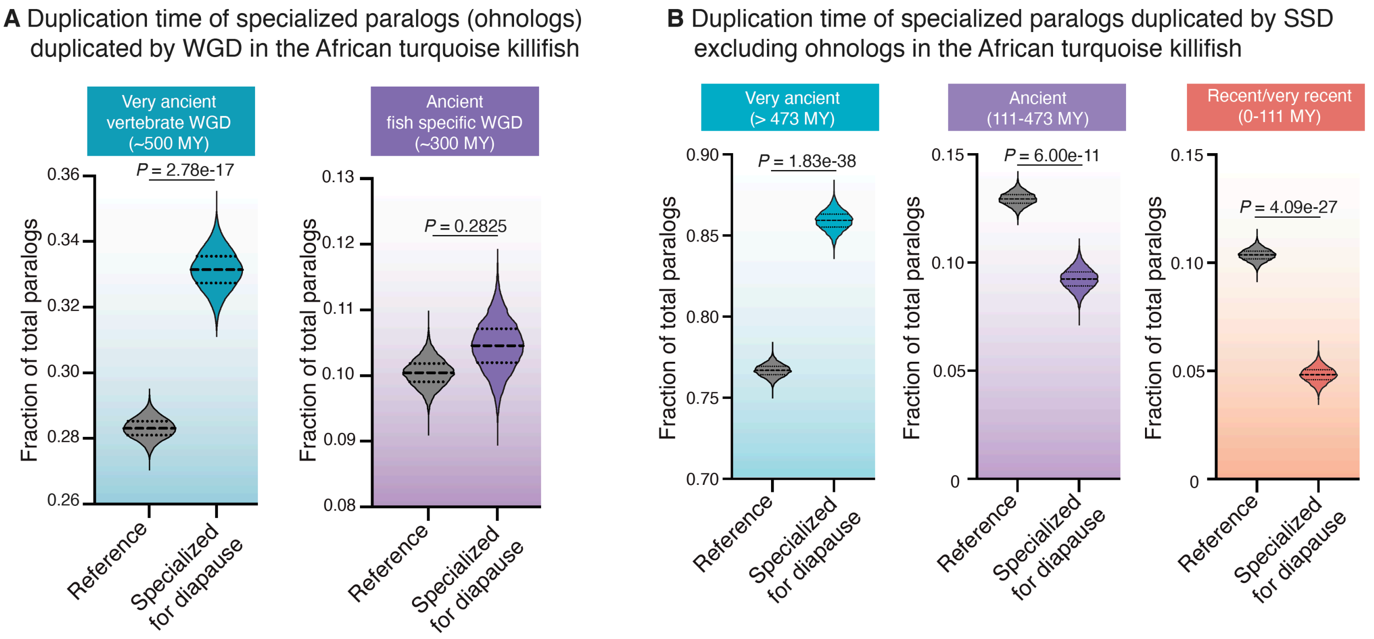

A Duplication time of specialized paralogs (ohnologs) duplicated by WGD in the African turquoise killifish
Very ancient vertebrate WGD (~500 MY)
P = 2.78e-17
Fraction of total paralogs
0.36
0.34
0.32
0.30
0.28
0.26
Reference
Specialized for diapause
Ancient fish specific WGD (~300 MY)
P = 0.2825
0.13
0.12
0.11
0.10
0.09
0.08
B Duplication time of specialized paralogs duplicated by SSD excluding ohnologs in the African turquoise killifish
Very ancient (> 473 MY)
P = 1.83e-38
0.90
0.85
0.80
0.75
0.70
Ancient (111-473 MY)
P = 6.00e-11
0.15
0.10
0.05
0
Recent/very recent (0-111 MY)
P = 4.09e-27

**Figure S3. Specialization of paralogs duplicated by whole genome duplication or small-scale duplication in the African turquoise killifish.** (A) Fraction of total paralog pairs (ohnologs) within either the vertebrate ancestor Whole Genome Duplication (WGD) event (left) or the fish ancestor WGD event (right). The enrichment of diapause-specialized paralogs pairs within each bin is compared to genome-wide expectation (reference). Violin plots represent distribution of observed vs expected specialized paralog fractions generated through 10,000 bootstrapped random sampling. Median and quartiles are indicated by dashed lines. Compared to the reference, paralogs with specialization in diapause are enriched among genes from the vertebrate ancestral WGD event and depleted among genes from the fish ancestral WGD event respectively. *P*-values from Chi-square test. (B) Fraction of total paralog pairs within each of the very ancient (left), ancient (middle), and recent/very recent (right) binned categories from Small Scale Duplication (SSD), after excluding ohnologs. Violin plots represent distribution of observed vs expected specialized paralog fractions generated through 10,000 bootstrapped random sampling. Median and quartiles are indicated by dashed lines. The enrichment of diapause-specialized paralogs pairs within each bin is compared to genome-wide expectation (reference). Compared to the reference, paralogs with specialization in diapause are enriched among genes with very ancient SSD duplication times and depleted among genes with ancient and recent/very recent SSD duplication times respectively. *P*-values from Chi-square test.

Figure S4

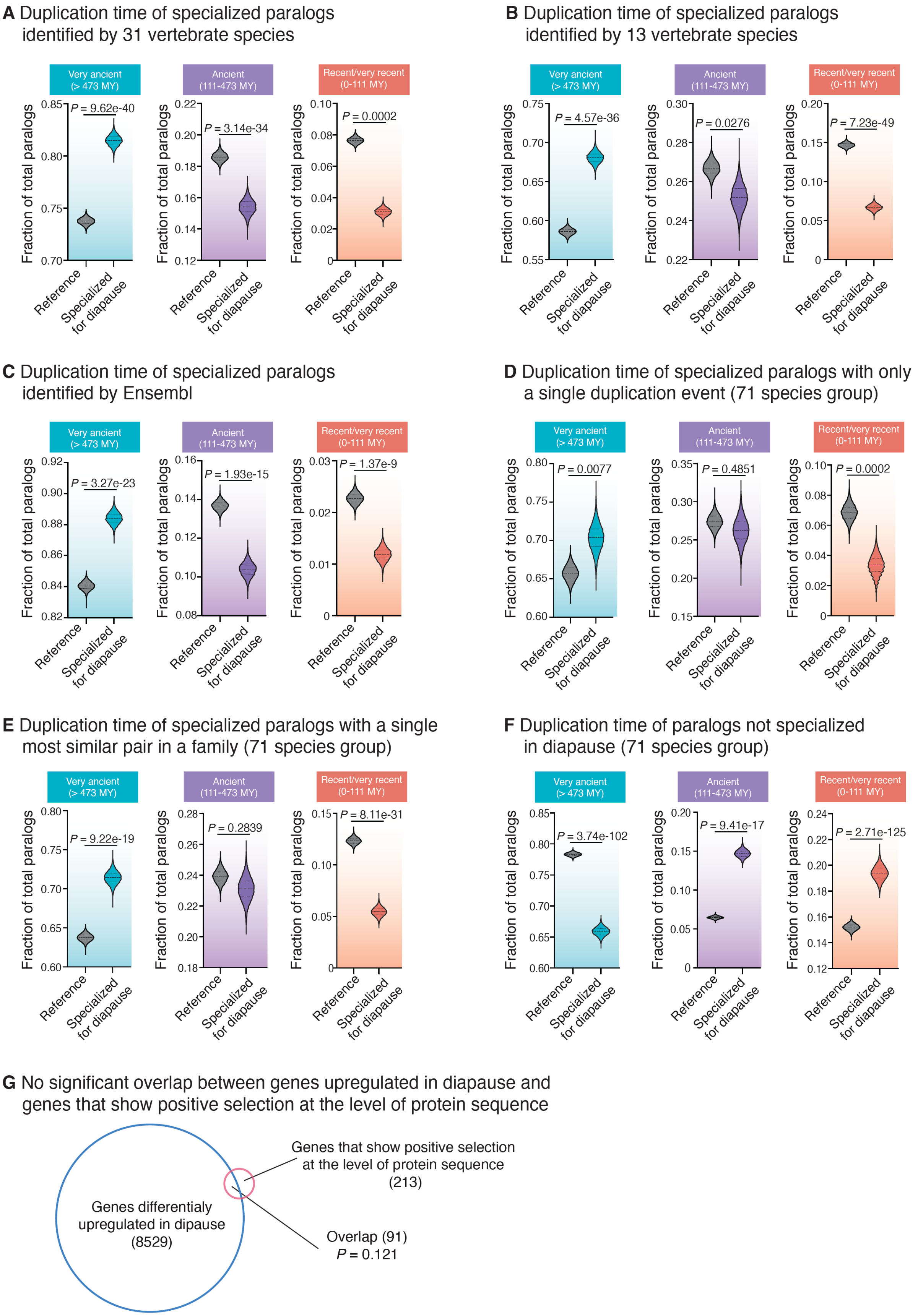

A Duplication time of specialized paralogs identified by 31 vertebrate species
Very ancient (> 473 MY)
Ancient (111-473 MY)
Recent/very recent (0-111 MY)
P = 9.62e-40
P = 3.14e-34
P = 0.0002
Fraction of total paralogs
Reference
Specialized for diapause
B Duplication time of specialized paralogs identified by 13 vertebrate species
P = 4.57e-36
P = 0.0276
P = 7.23e-49
C Duplication time of specialized paralogs identified by Ensembl
P = 3.27e-23
P = 1.93e-15
P = 1.37e-9
D Duplication time of specialized paralogs with only a single duplication event (71 species group)
P = 0.0077
P = 0.4851
P = 0.0002
E Duplication time of specialized paralogs with a single most similar pair in a family (71 species group)
P = 9.22e-19
P = 0.2839
P = 8.11e-31
F Duplication time of paralogs not specialized in diapause (71 species group)
P = 3.74e-102
P = 9.41e-17
P = 2.71e-125
G No significant overlap between genes upregulated in diapause and genes that show positive selection at the level of protein sequence
Genes that show positive selection at the level of protein sequence (213)
Genes differentialy upregulated in dipause (8529)
Overlap (91)
P = 0.121

**Figure S4. Specialization of paralogs in the African turquoise killifish with different paralog sources to assess robustness.** (A) Fraction of total paralog pairs within each of the very ancient (left), ancient (middle), and recent/very recent (right) binned categories using 31 rather than 71 species (fig. S2C). The enrichment of diapause-specialized paralogs pairs within each bin is compared to genome-wide expectation(reference). Violin plots represent distribution of observed vs expected specialized paralog fractions generated through 10,000 bootstrapped random sampling. Median and quartiles are indicated by dashed lines. Compared to the reference, paralogs with specialization in diapause are enriched among genes with very ancient duplication times and depleted among genes with ancient and recent or very recent duplication times respectively. *P*-values from Chi-square test. (B) Fraction of total paralog pairs within each of the very ancient (left), ancient (middle), and recent/very recent (right) binned categories using 13 rather than 71 species (fig. S2D). Violin plots represent distribution of observed vs expected specialized paralog fractions generated through 10,000 bootstrapped random sampling. Median and quartiles are indicated by dashed lines.The enrichment of diapause-specialized paralogs pairs within each bin is compared to genome-wide expectation (reference). Compared to the reference, paralogs with specialization in diapause are enriched among genes with very ancient duplication times and depleted among genes with ancient and recent or very recent duplication times respectively. *P*-values from Chi-square test. (C) Fraction of total paralog pairs within each of the very ancient (left), ancient (middle), and recent or very recent (right) binned categories using independent duplication time estimates from Ensembl rather than our pipeline (see methods). Violin plots represent distribution of observed vs expected specialized paralog fractions generated through 10,000 bootstrapped random sampling. The enrichment of diapause-specialized paralogs pairs within each bin is compared to genome-wide expectation (reference). Median and quartiles are indicated by dashed lines. Compared to the reference, paralogs with specialization in diapause are enriched among genes with very ancient duplication times and depleted among genes with ancient and recent or very recent duplication times, respectively. *P*-values from Chi-square test. (D) Fraction of total paralog pairs within each of the very ancient (left), ancient (middle), and recent/very recent (right) binned categories. Violin plots represent distribution of observed vs expected specialized paralog fractions generated through 10,000 bootstrapped random sampling. Median and quartiles are indicated by dashed lines. Only paralogs that have experienced a single duplication event were included in this analysis. The enrichment of diapause-specialized paralogs pairs within each bin is compared to genome-wide expectation (reference). Compared to the reference, paralogs with specialization in diapause are enriched among genes with very ancient duplication times and depleted among genes with ancient and recent or very recent duplication times respectively, indicating that our results are not affected by gene family size. *P*-values from Chi-square test. (E) Fraction of total paralog pairs within each of the very ancient (left), ancient (middle), and recent/very recent (right) binned categories. For each gene family only a single paralog pair with highest similarity was used (e.g. in the following tree, (A, (B, C)), only pair B-C would be kept while A-B and A-C would be discarded). The enrichment of diapause-specialized paralogs pairs within each bin is compared to genome-wide expectation (reference). Compared to the reference, paralogs with specialization in diapause are enriched among genes with very ancient duplication times and depleted among genes with ancient and recent or very recent duplication times respectively, indicating that our results are not affected by gene family size. (F) Fraction of total paralog pairs within each of the very ancient (left), ancient (middle), and recent/very recent (right) binned categories. Violin plots represent distribution of observed vs expected specialized paralog fractions generated through 10,000 bootstrapped random sampling. Median and quartiles are indicated by dashed lines. The enrichment of non-diapause-specialized paralogs pairs within each bin is compared to genome-wide expectation (reference). Compared to the reference, paralogs with no specialization in diapause are depleted among genes with very ancient duplication times and enriched among genes with ancient and recent or very recent duplication times respectively, suggesting that our results are specific to diapause-specialized paralogs. (G) Overlap between genes upregulated during diapause in the African turquoise killifish (blue circle) and genes from the African turquoise killifish that showed a signature of positive selection at the level of protein sequence (red circle). The overlap between the two categories is not significant ($P = 0.121$, hypergeometric test).

Figure S5

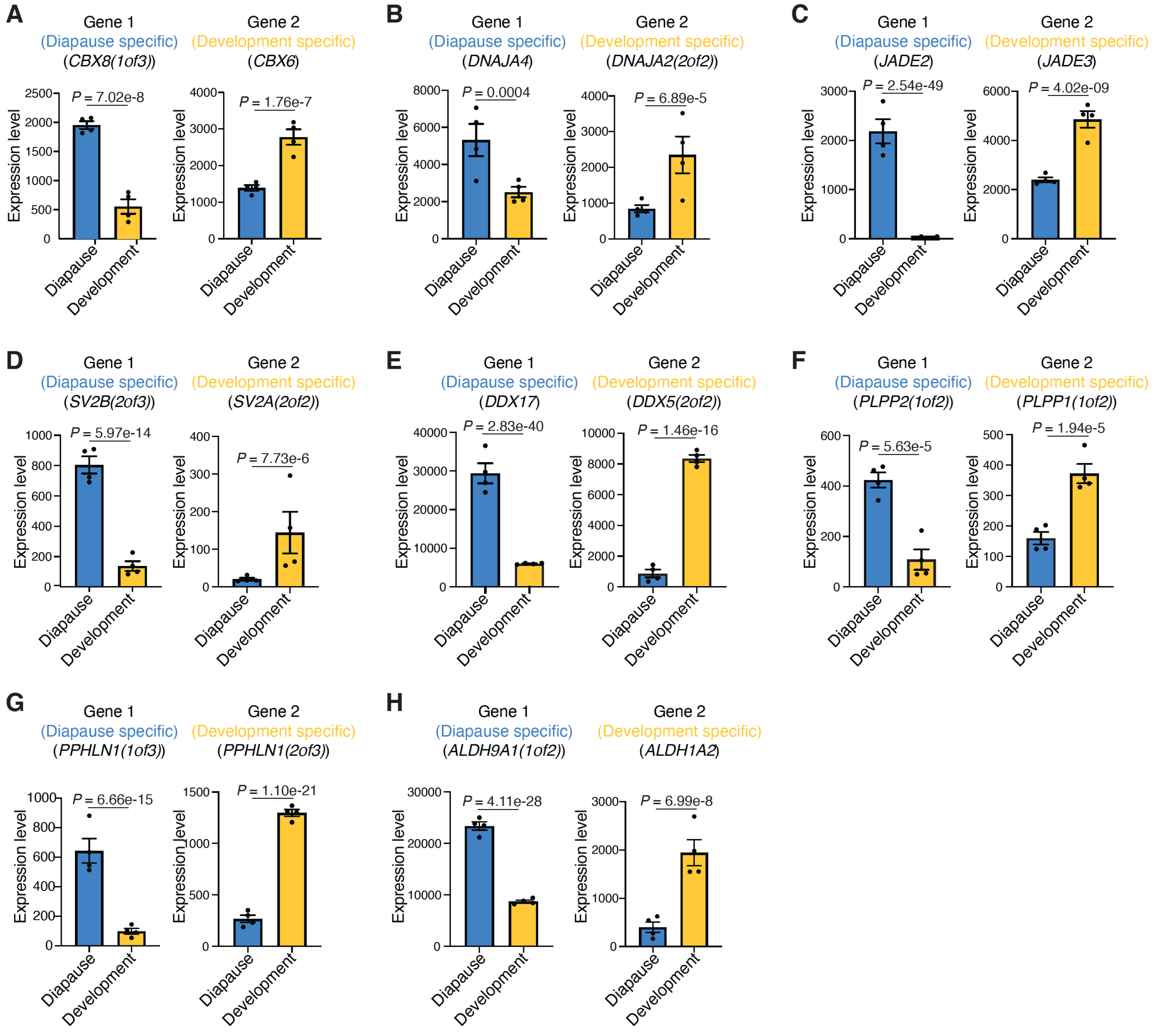
A
Gene 1
(Diapause specific)
(CBX8(1of3))
P = 7.02e-8
Gene 2
(Development specific)
(CBX6)
P = 1.76e-7
B
Gene 1
(Diapause specific)
(DNAJA4)
P = 0.0004
Gene 2
(Development specific)
(DNAJA2(2of2))
P = 6.89e-5
C
Gene 1
(Diapause specific)
(JADE2)
P = 2.54e-49
Gene 2
(Development specific)
(JADE3)
P = 4.02e-09
D
Gene 1
(Diapause specific)
(SV2B(2of3))
P = 5.97e-14
Gene 2
(Development specific)
(SV2A(2of2))
P = 7.73e-6
E
Gene 1
(Diapause specific)
(DDX17)
P = 2.83e-40
Gene 2
(Development specific)
(DDX5(2of2))
P = 1.46e-16
F
Gene 1
(Diapause specific)
(PLPP2(1of2))
P = 5.63e-5
Gene 2
(Development specific)
(PLPP1(1of2))
P = 1.94e-5
G
Gene 1
(Diapause specific)
(PPHLN1(1of3))
P = 6.66e-15
Gene 2
(Development specific)
(PPHLN1(2of3))
P = 1.10e-21
H
Gene 1
(Diapause specific)
(ALDH9A1(1of2))
P = 4.11e-28
Gene 2
(Development specific)
(ALDH1A2)
P = 6.99e-8
Expression level
Diapause
Development

**Figure S5. Additional examples of diapause-specialized paralogs in South American killifish.** (A-H) Examples of paralog gene pairs, with specialized expression of gene 1 in diapause (blue) and gene 2 in development (yellow) in South American killifish (*Austrofundulus limnaeus*). Displayed gene names are the assigned name of relevant ortholog in African turquoise killifish for comparison. Bars represent mean expression level (normalized DESeq2 count) across replicates in diapause or development state. Dots show normalized DESeq2 counts in each replicate. Error bar is standard error of mean (SEM). *P*-values from DESeq2 Wald test.

Figure S6

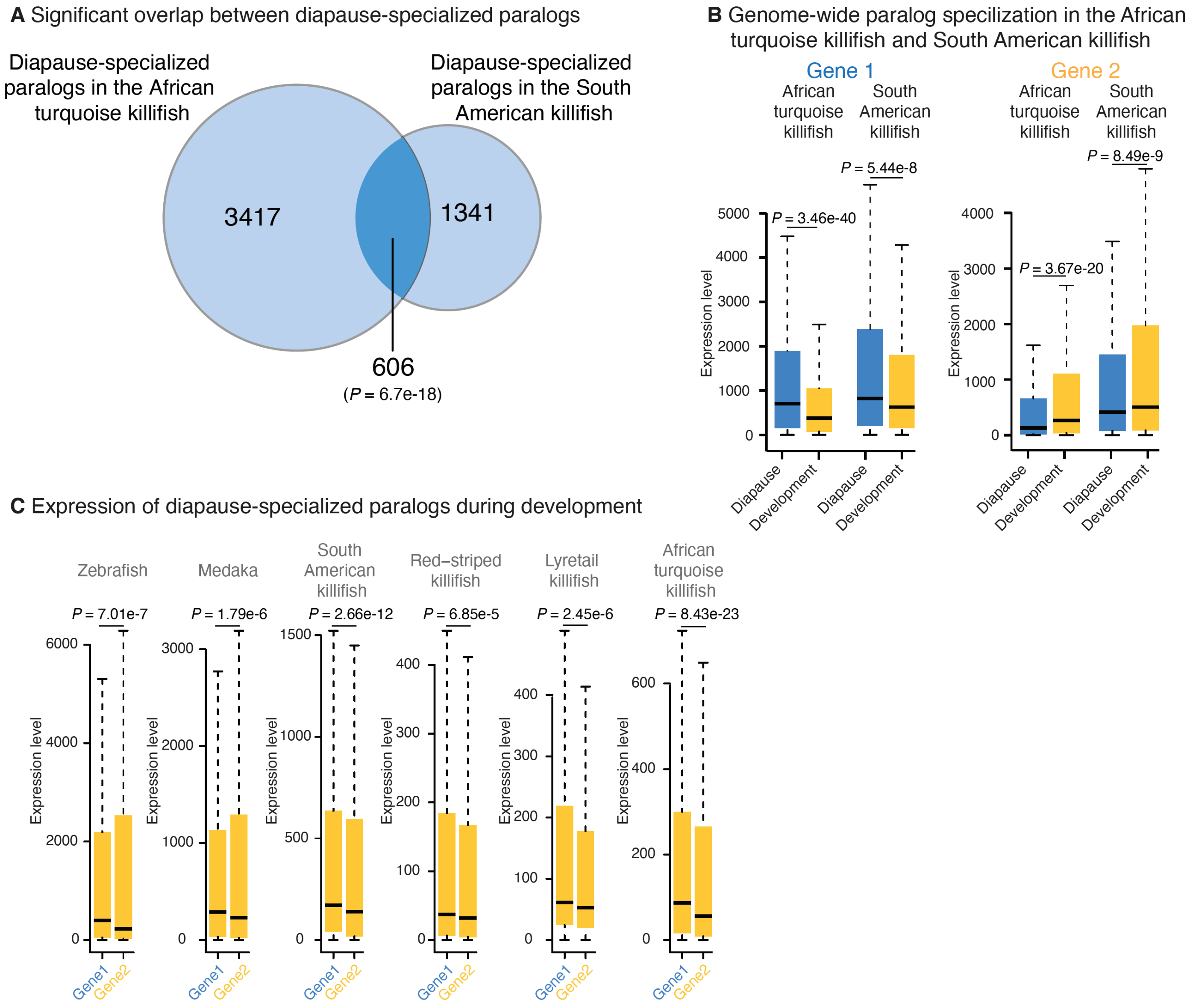
A Significant overlap between diapause-specialized paralogs
Diapause-specialized paralogs in the African turquoise killifish
Diapause-specialized paralogs in the South American killifish
3417
1341
606
(P = 6.7e-18)
B Genome-wide paralog specilization in the African turquoise killifish and South American killifish
Gene 1
Gene 2
African turquoise killifish
South American killifish
P = 3.46e-40
P = 5.44e-8
P = 3.67e-20
P = 8.49e-9
Expression level
Diapause
Development
C Expression of diapause-specialized paralogs during development
Zebrafish
Medaka
South American killifish
Red–striped killifish
Lyretail killifish
African turquoise killifish
P = 7.01e-7
P = 1.79e-6
P = 2.66e-12
P = 6.85e-5
P = 2.45e-6
P = 8.43e-23
Expression level
Gene1
Gene2

**Figure S6. Comparison of diapause gene expression and specialized paralogs in African turquoise killifish and South American killifish.** (A) Overlap between paralog pairs specialized for diapause in both African turquoise killifish and South American killifish. Paralog pairs were included only if both the genes in the pairs were orthologous to each other with the same duplication time. There is a significant overlap of these pairs between the two killifish species ($P$ = 6.7e-18, Hypergeometric test). (B) Comparison of diapause-specialized paralogs identified in the African turquoise killifish to their orthologs in the South American killifish. Both the diapause-specialized genes (*Gene 1* cohort; left panel) and the development-specialized gene (*Gene 2* cohort; right panel) exhibit the same expression pattern genome-wide in both the fish species. These expression differences were significant in both African turquoise killifish and South American killifish. $P$-values from Kolmogorov-Smirnov test. (C) Expression of African turquoise killifish diapause-specialized paralogs and their one-to-one orthologs in other species. Expression was evaluated during Pre-Diapause time point and $P$-values were calculated using Kolmogorov-Smirnov test. In all species evaluated, the expression pattern was similar at the comparable pre-diapause developmental time point with diapause-specific gene (Gene 1) always the more highly expressed during the pre-diapause developmental time point. This expression asymmetry is a known property of paralogs (*24*).

Figure S7

**A** Identification of diapause-specific ATAC-seq peaks in African turquoise killifish

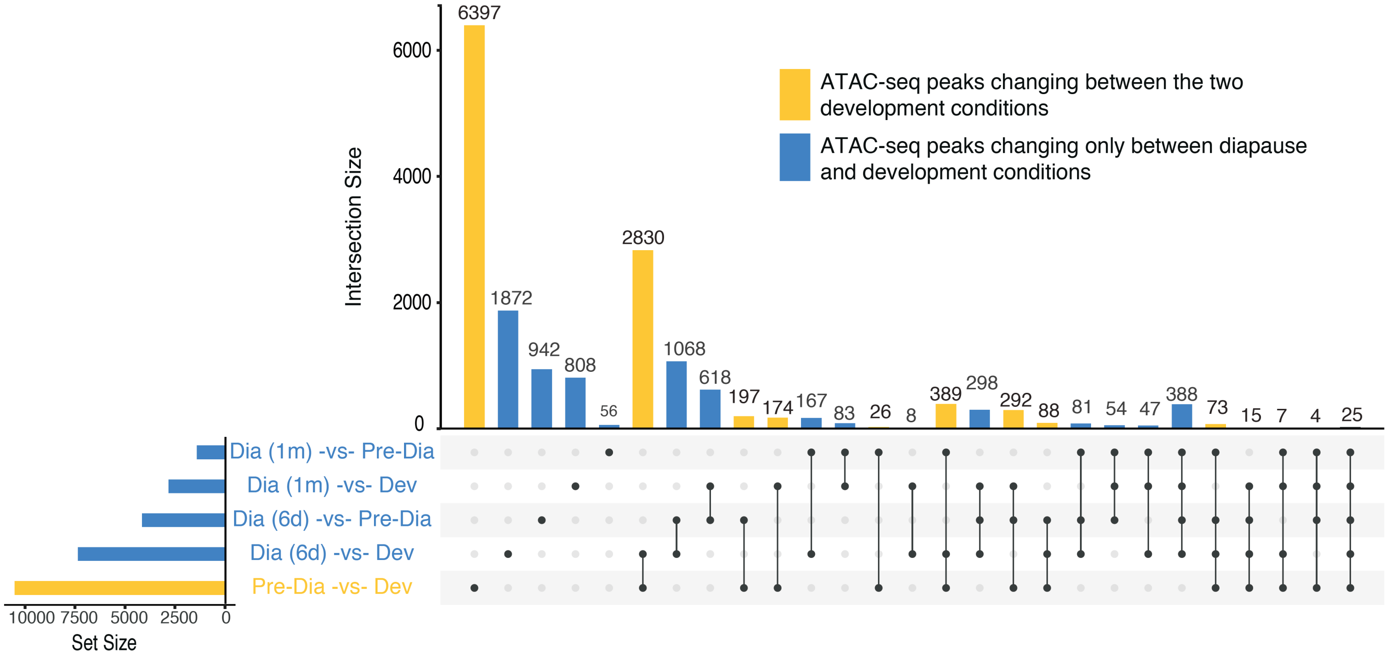


**B** Feature distribution of differential ATAC-seq peaks in diapause in African turquoise killifish

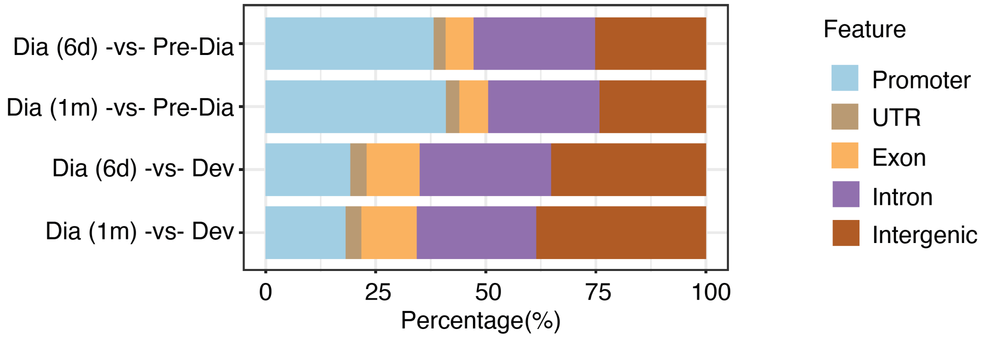


**C** Correlation between RNA-seq and ATAC-seq in diapause in African turquoise killifish

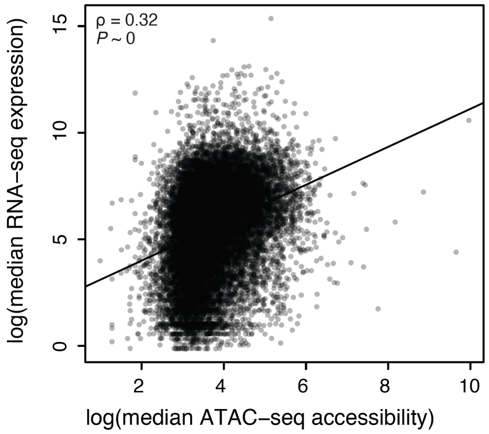


**D** Correlation between RNA-seq and ATAC-seq in development in African turquoise killifish

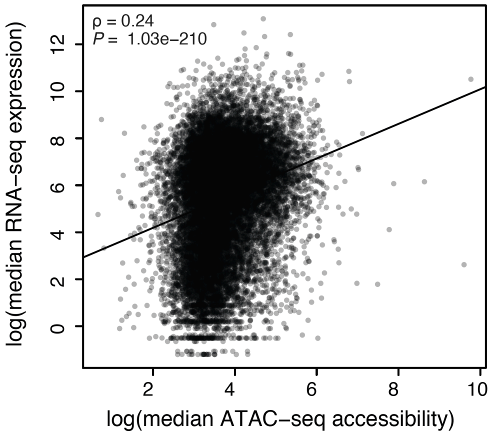

**Figure S7. Characterization of ATAC-seq datasets in the African turquoise killifish and other species.** (A) Upset plot depicting differentially accessible chromatin regions (ATAC-seq peaks) between the consensus peak set for each biological timepoint surveyed in the African turquoise killifish. The final set of differentially accessible chromatin regions used for analysis is comprised of all intersections containing peaks that only change between diapause and development conditions (blue histogram bins) while those that include a change between developmental timepoints were excluded (yellow histogram bins). (B) Percentage breakdowns of included diapause-specific, differentially accessible chromatin regions by genome feature in the African turquoise killifish. Feature categories (Promoter, UTR, Exon, Intron, and Intergenic) were made by consolidating more specific sub-feature categories provided by the DESeq2 pipeline. (C) Correlation plot between the median gene expression (RNA-seq) and the median chromatin accessibility (ATAC-seq) for all genes during diapause in African turquoise killifish ($P \sim 0$, Spearman correlation coefficient = 0.32). (D) Correlation plot between the median gene expression (RNA-seq) and the median chromatin accessibility (ATAC-seq) for all genes during development in the African turquoise killifish ($P$ = 1.03e-210, Spearman correlation coefficient = 0.24)

Figure S8

**A** ATAC-seq library Transcription Start Site (TSS) enrichment in the African turquoise killifish and other killifish species

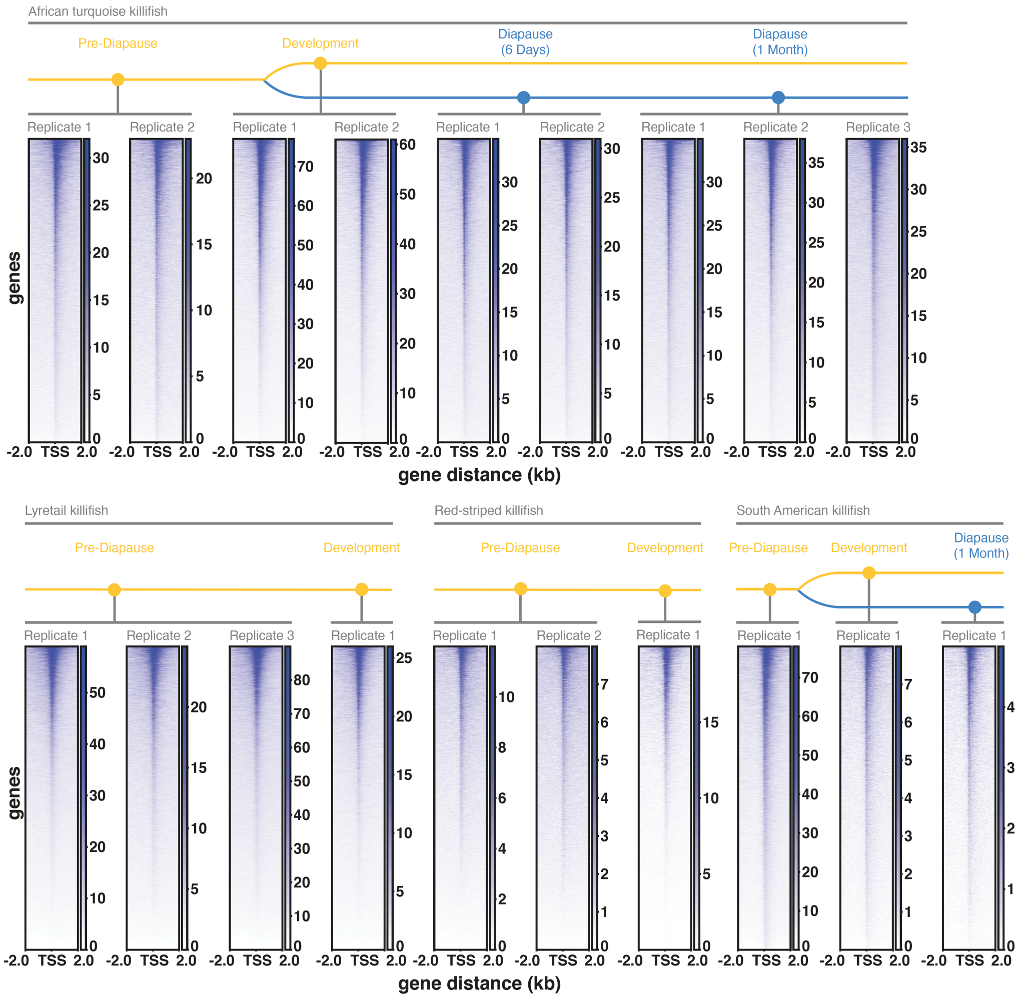


**B** ATAC-seq library nucleosome banding in the African turquoise killifish and other killifish species

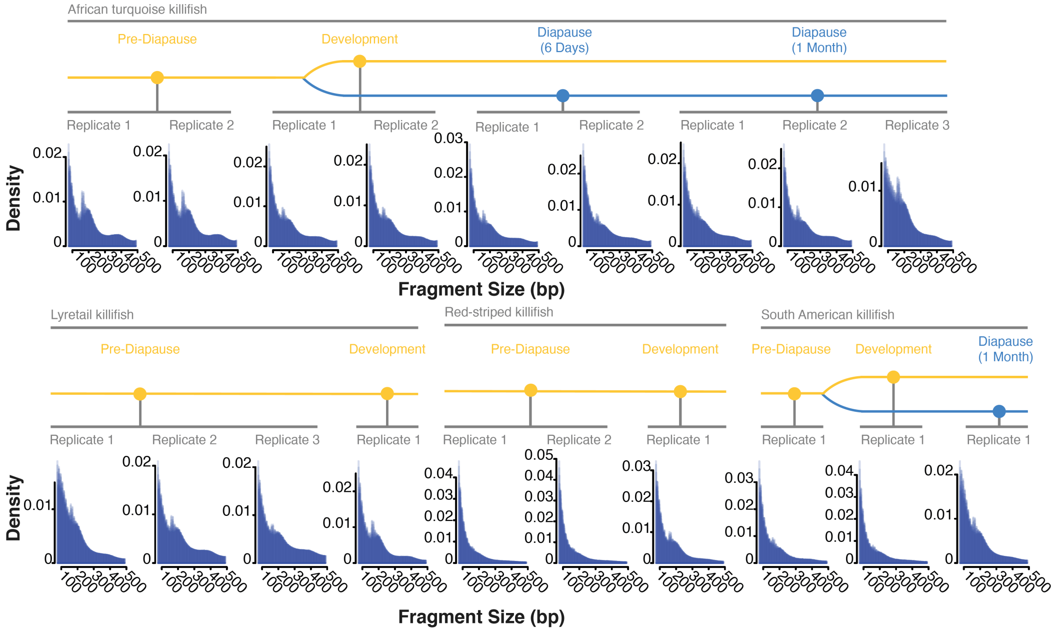

**Figure S8. ATAC-seq library quality metrics in African turquoise killifish and other species.** (A) TSS read enrichment compared to neighboring 2kb regions for each ATAC-seq library generated. An enrichment of accessibility signal at TSS indicates good quality. (B) Nucleosome banding patten displaying the presence/absence and intensity of the mono-, di-, and tri-nucleosome bands for each ATAC-seq library.

Figure S9

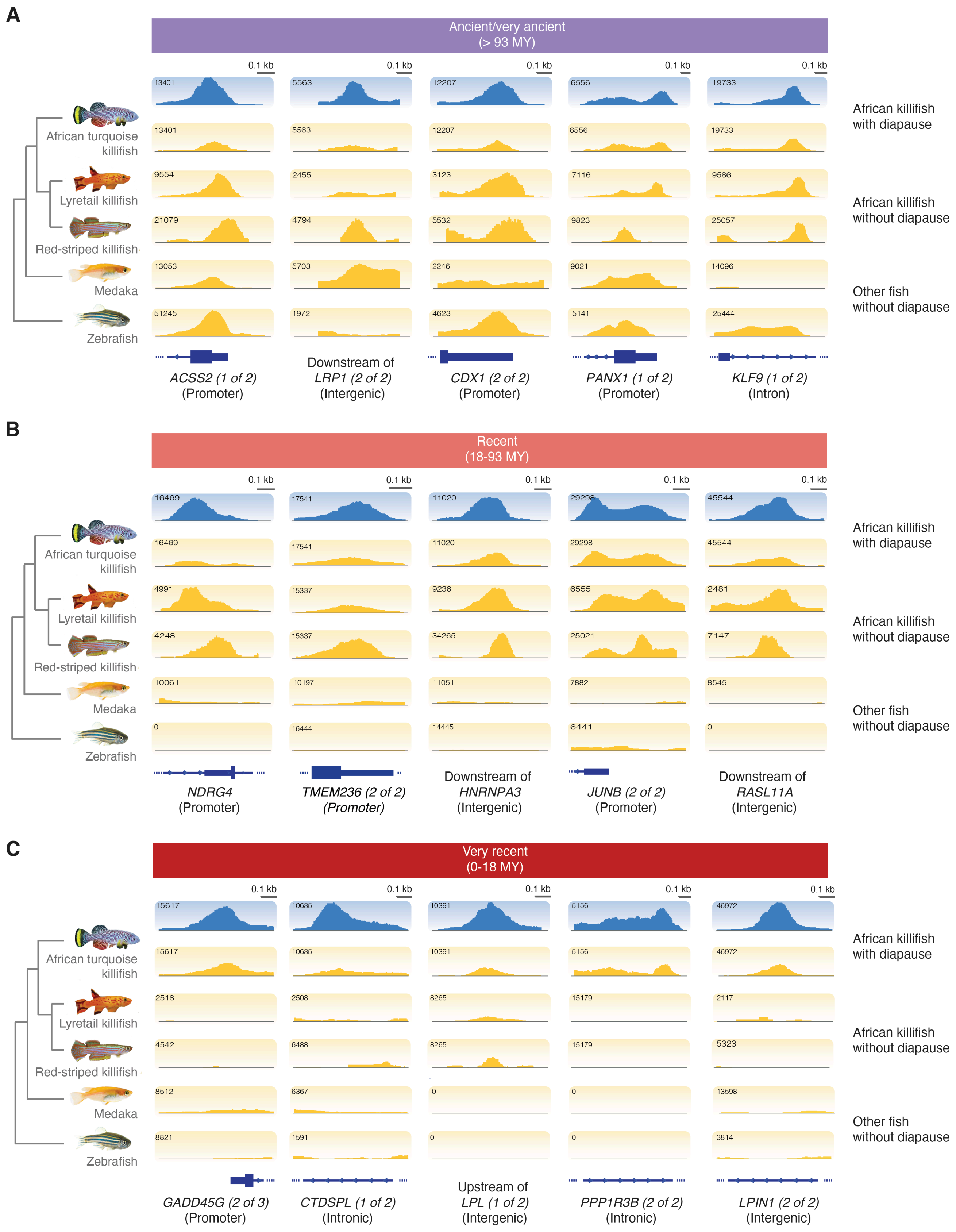

A
Ancient/very ancient
(> 93 MY)
African turquoise killifish
Lyretail killifish
Red-striped killifish
Medaka
Zebrafish
African killifish with diapause
African killifish without diapause
Other fish without diapause
ACSS2 (1 of 2) (Promoter)
Downstream of LRP1 (2 of 2) (Intergenic)
CDX1 (2 of 2) (Promoter)
PANX1 (1 of 2) (Promoter)
KLF9 (1 of 2) (Intron)
B
Recent
(18-93 MY)
NDRG4 (Promoter)
TMEM236 (2 of 2) (Promoter)
Downstream of HNRNPA3 (Intergenic)
JUNB (2 of 2) (Promoter)
Downstream of RASL11A (Intergenic)
C
Very recent
(0-18 MY)
GADD45G (2 of 3) (Promoter)
CTDSPL (1 of 2) (Intronic)
Upstream of LPL (1 of 2) (Intergenic)
PPP1R3B (2 of 2) (Intronic)
LPIN1 (2 of 2) (Intergenic)

**Figure S9. Additional examples of diapause-accessible (differential) ATAC-seq peaks and their cross-species conservation.** IGV visualization tracks of chromatin accessibility for representative ATAC-seq peaks across African turquoise, lyretail, and red-striped killifish in addition to medaka and zebrafish from RPKM-normalized reads summed across replicates and biological timepoints (e.g., diapause and development separately) to obtain single tracks for each species. The tree of species represents labeling each track displays the evolutionary relationship between each evaluated species. Peaks displayed showcase the three conservation categories evaluated. (A) Conserved chromatin accessibility across all species (ancient/very ancient). (B) Conserved chromatin accessibility exclusive to surveyed killifish species (recent). (C) Chromatin accessibility exclusive to the African turquoise killifish (very recent). Each region is labeled as one of the three types of genomic features on which peaks were evaluated: promoters, introns, and intergenic regions. Peaks located in proximity to *RASL11A* (B, column 5) and *GADD45G* (C, column 1) are at specialized paralogs (fig. S1G and S1H, respectively).

#### Figure S10

**A** Genome-wide assessment of sequence and chromatin conservation

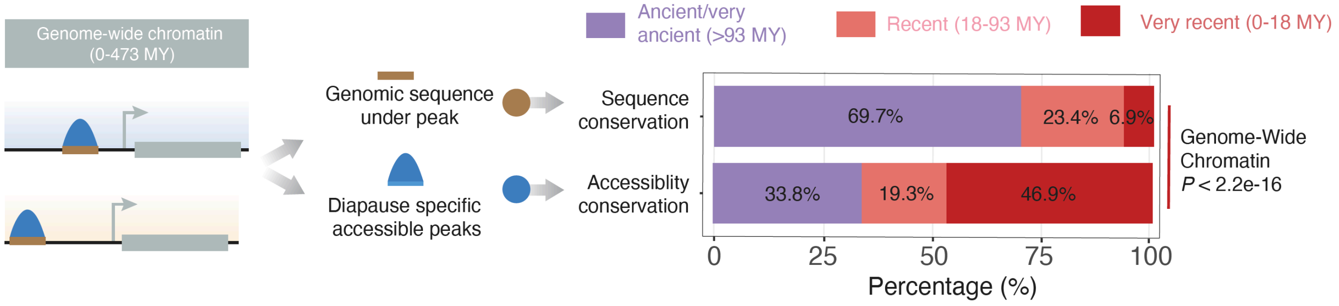


**B** Assessment of diapause-accessible chromatin at ancient paralogs by genomic feature

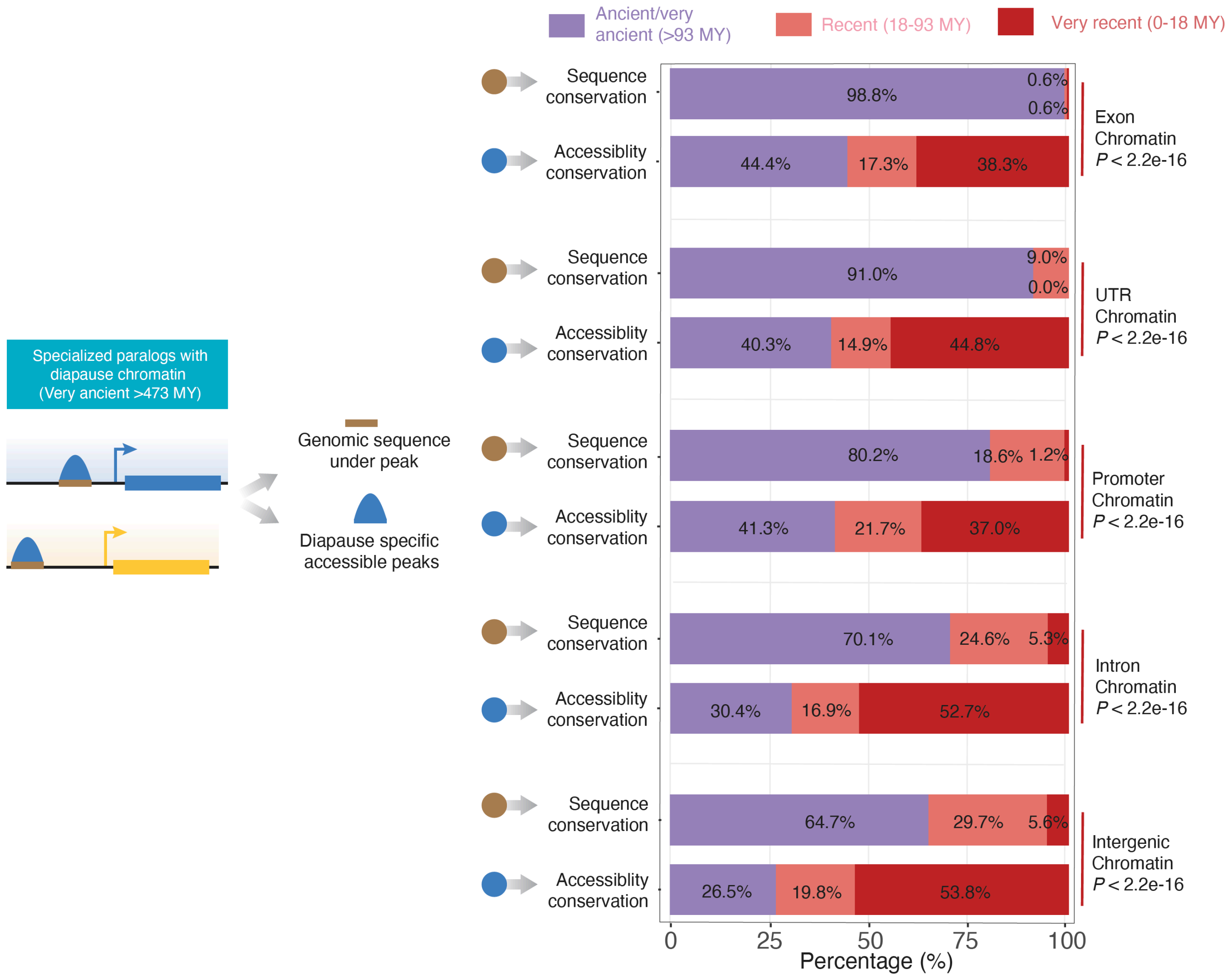

**Figure S10. Genomic sequence and chromatin accessibility conservation breakdown.** (A) Conservation analysis of genomic sequence and chromatin accessibility genome-wide for all the significant diapause specific chromatin peaks (see Fig. 3E for paralog specific result). Left panel: Schematic of the analysis. Right panel: Percentage (e.g. conservation) of alignable regions containing diapause-specific chromatin accessibility (upper) and the conservation of diapause-specific chromatin accessibility (lower) genome-wide (B) Conservation analysis of genomic sequence and chromatin accessibility at very ancient paralogs with specialization in diapause vs. development delineated by genomic feature in order of decreasing conservation: accessible chromatin in exons (upper pair), untranslated regions (UTRs) (upper-middle pair), promoters (middle pair), introns (middle-lower pair), and intergenic regions (lower pair). Left panel: Schematic of the analysis. Right panel: Percentage (e.g. conservation) of alignable regions containing diapause-specific chromatin accessibility (upper) and the conservation of diapause-specific chromatin accessibility (lower) near specialized ancient paralogs.

Figure S11

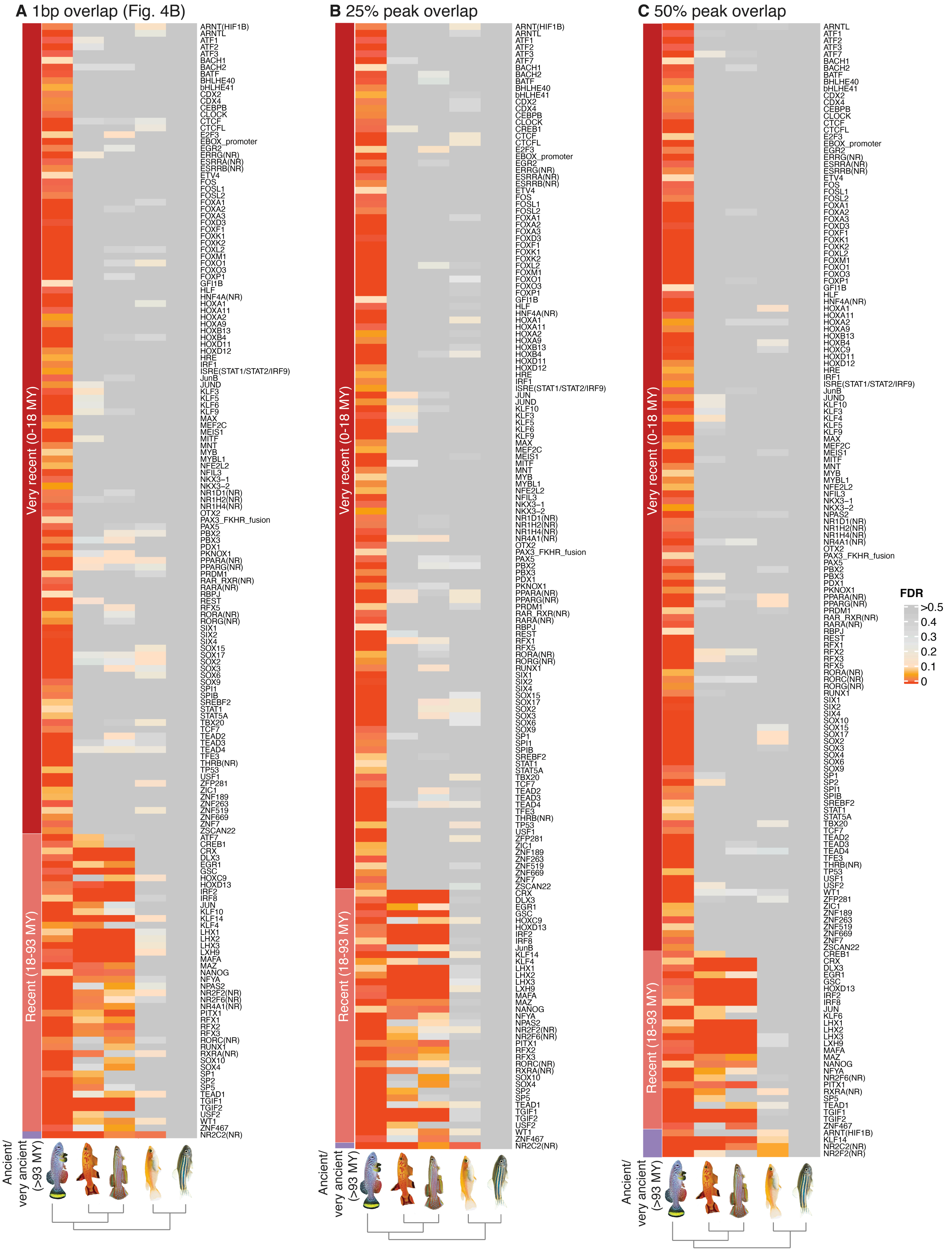
A 1bp overlap (Fig. 4B)
B 25% peak overlap
C 50% peak overlap
Very recent (0-18 MY)
Recent (18-93 MY)
Ancient/ very ancient (>93 MY)
FDR
>0.5
0.4
0.3
0.2
0.1
0

**Figure S11. Transcription-factor binding motifs enrichment across species using various conservation cutoffs.** Conservation in other fish species of transcription-factor binding motifs enriched in diapause-specific chromatin accessible regions in the African turquoise killifish. (A) For accessible chromatin regions (ATAC-seq peaks) in other species with at least a single base pair overlap with a diapause-specific chromatin accessible regions in the African turquoise killifish. (B) For accessible chromatin regions (ATAC-seq peaks) in other species with at least 25% peak overlap with a diapause-specific chromatin accessible regions in the African turquoise killifish. (C) For accessible chromatin regions (ATAC-seq peaks) in other species with at least 50% peak overlap with a diapause-specific chromatin accessible regions in the African turquoise killifish. The majority of diapause-specific motifs are very recent (i.e. specific to African turquoise killifish) and are not enriched in killifish species without diapause or outgroup species with all the three criteria.

Figure S12

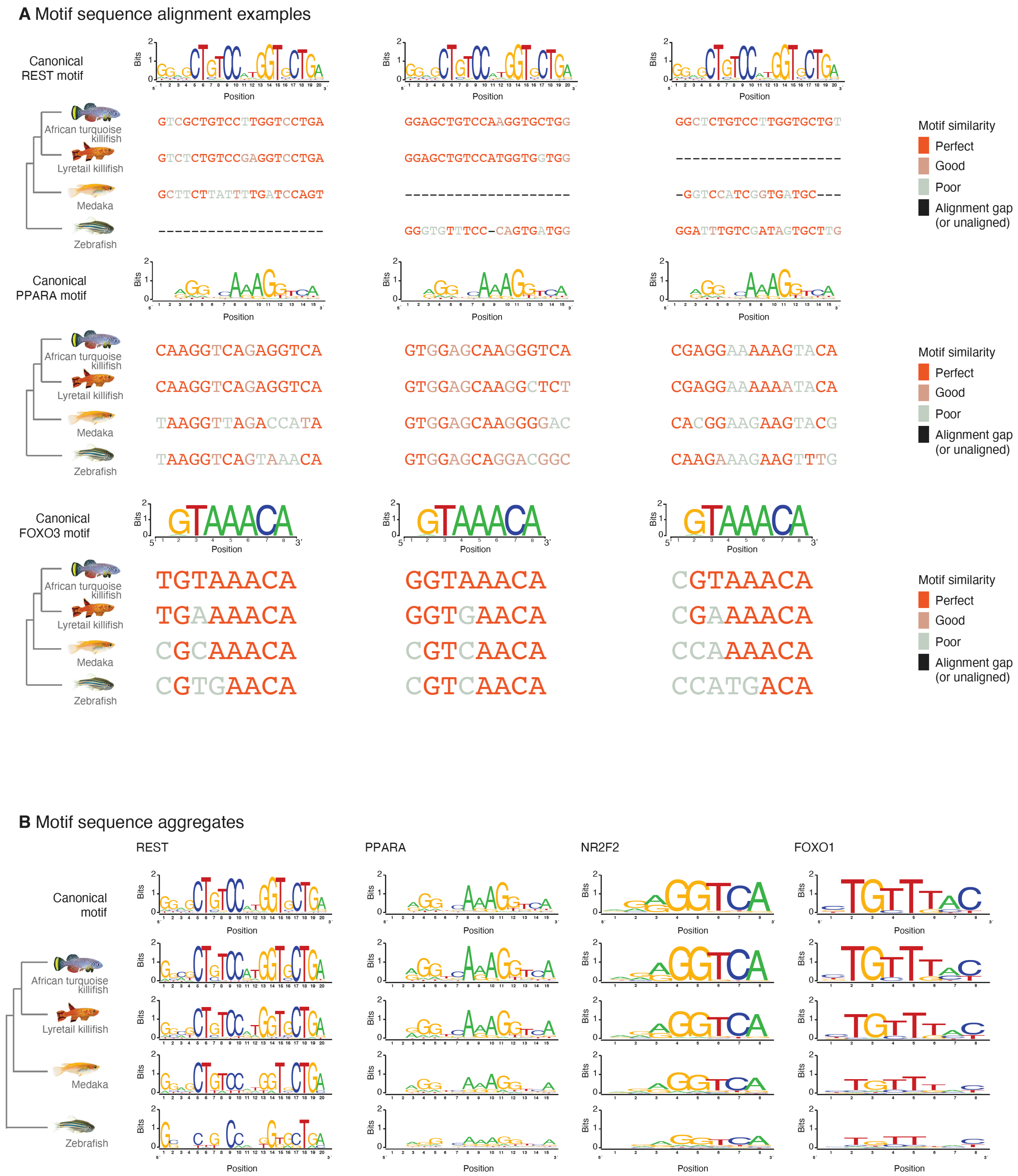
A Motif sequence alignment examples
Canonical REST motif
Canonical PPARA motif
Canonical FOXO3 motif
Bits
Position
African turquoise killifish
Lyretail killifish
Medaka
Zebrafish
GTCGCTGTCCTTGGTCCTGA
GTCTCTGTCCGAGGTCCTGA
GCTTCTTATTTTGATCCAGT
--------------------
GGAGCTGTCCAAGGTGCTGG
GGAGCTGTCCATGGTGGTGG
--------------------
GGGTGTTTCC-CAGTGATGG
GGCTCTGTCCTTGGTGCTGT
--------------------
-GGTCCATCGGTGATGC---
GGATTTGTCGATAGTGCTTG
CAAGGTCAGAGGTCA
CAAGGTCAGAGGTCA
TAAGGTTAGACCATA
TAAGGTCAGTAAACA
GTGGAGCAAGGGTCA
GTGGAGCAAGGCTCT
GTGGAGCAAGGGGAC
GTGGAGCAGGACGGC
CGAGGAAAAAGTACA
CGAGGAAAAAATACA
CACGGAAGAAGTACG
CAAGAAAGAAGTTTG
TGTAAACA
TGAAAACA
CGCAAACA
CGTGAACA
GGTAAACA
GGTGAACA
CGTCAACA
CGTCAACA
CGTAAACA
CGAAAACA
CCAAAACA
CCATGACA
Motif similarity
Perfect
Good
Poor
Alignment gap (or unaligned)
B Motif sequence aggregates
REST
PPARA
NR2F2
FOXO1
Canonical motif

**Figure S12. Additional examples and aggregates from motif evolution analysis.** (A) Representative examples of REST (upper), PPARA (middle), and FOXO3 (lower) transcription factor binding sites in African turquoise and the aligned regions in other evaluated fish species. Aligned sequences colored in accordance with their closeness-of-fit to the information content of HOMER-produced consensus motif logo (top track). Only a single sequence is provided for both lyretail killifish and red-striped killifish as they are aligned to the same draft genome. (B) Aggregated informational content (bits) across all REST (left), PPARA (left-center), NR2F2 (right-center), and FOXO1 (right) transcription factor binding sites in diapause-accessible (differential) chromatin and aligned regions in other species regardless of accessibility status. The canonical motif logos are provided for comparison (upper logo). During sequence aggregation gaps were removed along with sequence aligned to gaps (i.e., exact base pair to base pair alignment with the African turquoise killifish was used).

Figure S13

**A** Schematic of ancestral reconstruction

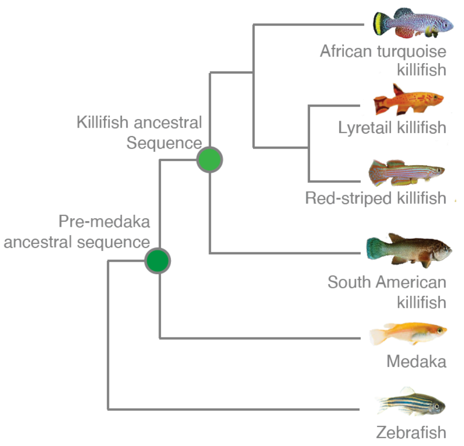


**B** Overlap of positive selection at regulatory regions using multiple ancestral sets

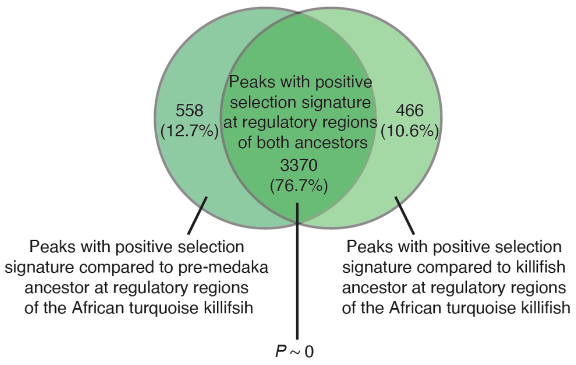


**C** Motif enrichment of peaks with positive selection signature at regulatory regions

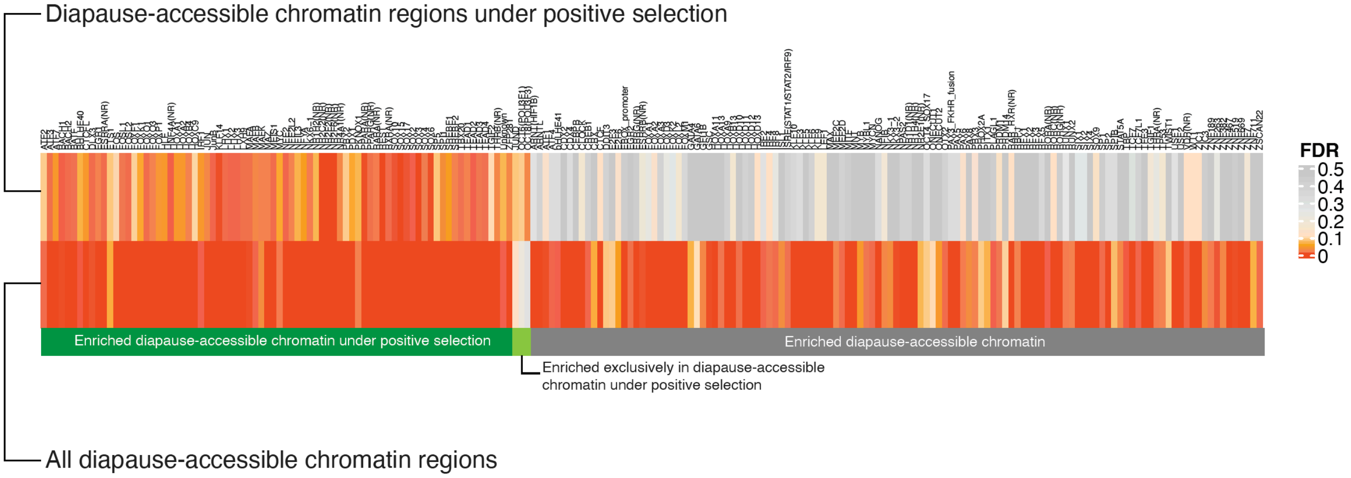

**Figure S13. Positive selection analysis and motif enrichment in accessible chromatin regions.** (A) Schematic tree showing the evolutionary timing of inferred ancestral sequences used for positive selection analysis on accessible chromatin regions (*99*). The green dots represent the inferred pre-medaka and killifish common ancestral sequences. The ancestral sequences were constructed using aligned sequences from each species in the tree with the site of green dots delineating the branches of the phylogeny classified as in-group and out-group respectively (see methods). (B) The overlap between peaks with a positive selection signature as calculated using the inferred pre-medaka (left) and killifish (right) ancestral sequence respectively. The overlap between the two was significant ($P \sim 0$, hypergeometric test). We used the union of the two sets for determining the positive selection signature overlap with diapause-accessible (differential) chromatin near ancient, specialized paralogs (Fig. 4G). (C) Enrichment of transcription factor binding motifs among diapause-accessible chromatin peaks near ancient, specialized paralogs with a positive selection signature. Motifs such as REST, FOXO and PPAR are significantly enriched in the positively selected chromatin regions.

Figure S14 Examples of transposable elements overlapping with TF binding sites

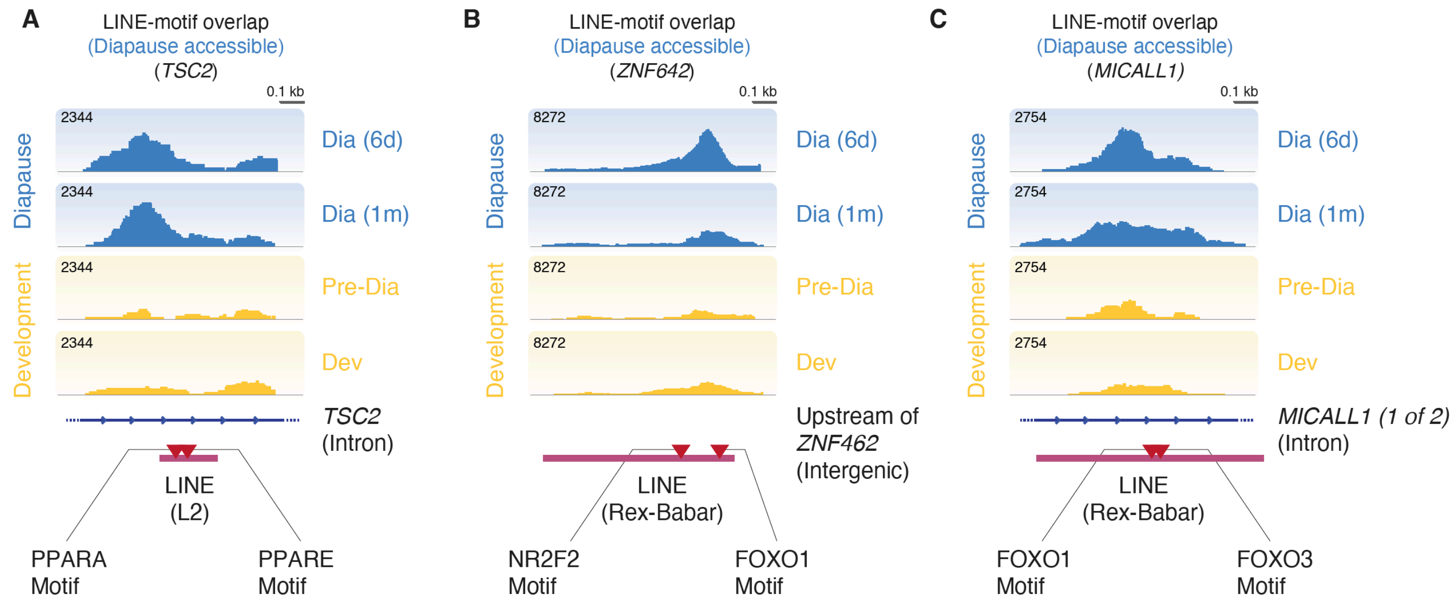

**Figure S14. Additional examples of TE overlapping transcription factor binding sites in African turquoise killifish.** (A-C) Genome browser (IGV) chromatin accessibility track of representative example peaks containing a TE-embedded transcription factor binding site. ATAC-seq library timepoints for both diapause (6 days and 1 month post entry) and development (Pre-Diapause and Development) are represented by replicated-summed, RPKM-normalized tracks. Blue lines at the bottom represent the genomic architecture (intron) at the region and magenta boxes display the location and size of TEs in the region with the motif binding sequence marked (red triangle).

Figure S15

**A** Significantly different lipid classes in diapause

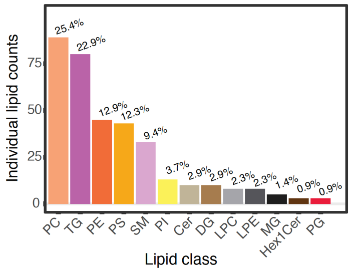


**B** Triglycerides metabolism enzymes are upregulated, and many regulators are differentially expressed in diapause

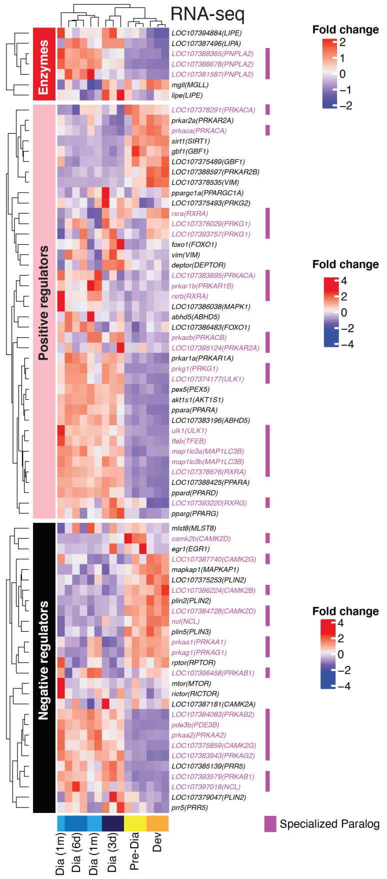


**C** Significant class-specific triglyceride content fold-change between killifish species

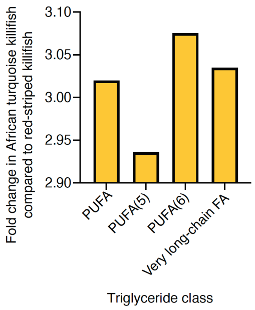


**D** Lipid abundance fold changes between African turquoise killifish time course and red-striped killifish

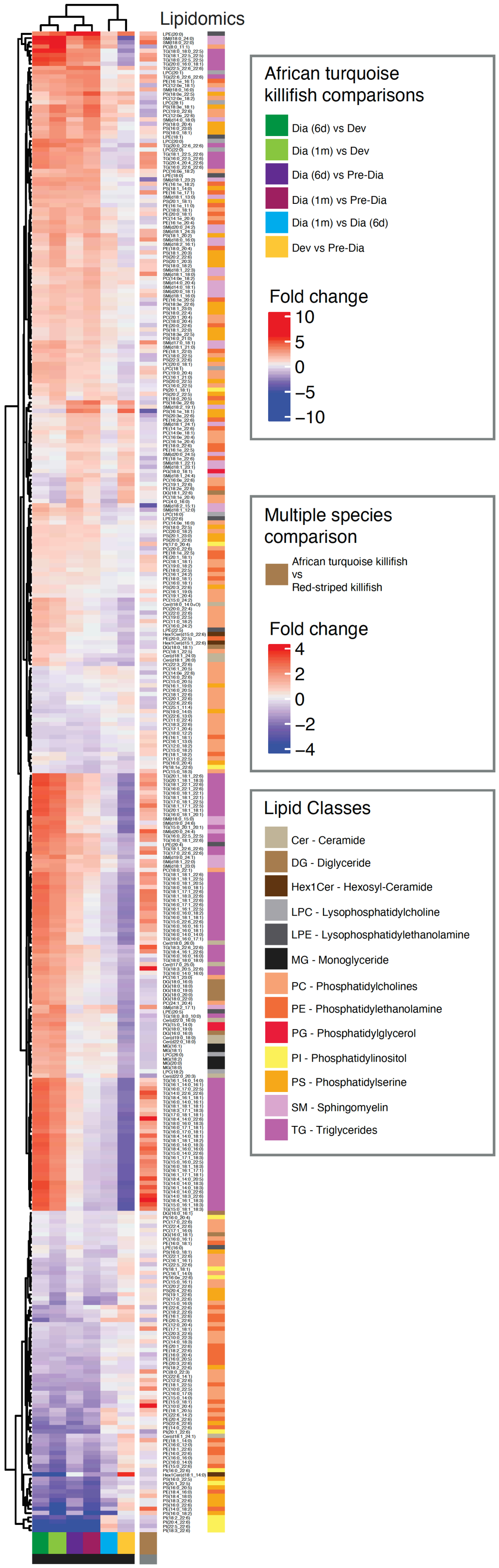

**Figure S15. Analysis of class level triglycerides (TG) in the African turquoise killifish and lyretail killifish.** (A) Bar graph representing the number of diapause-specific differential lipids in each lipid class. Phosphatidylcholines (PC) and Triglycerides (TG) constitute most of the differential lipids that change in diapause. PE, Phosphatidylethanolamine; PS, Phosphatidylserine; SM, Sphingomyelin; PI, Phosphatidylinositol; Cer, Ceramide; DG, Diglyceride; LPC, Lysophosphatidylcholine; LPE, Lysophosphatidylethanolamine; MG, Monoacylglyceride; Hex1Cer, Hexosyl-Ceramide; PG, Phosphatidylglycerol. (B) RNA-seq expression levels of the genes involved in triglyceride metabolism divided by their functions: enzymes (upper heatmap), positive regulators of triglyceride metabolism (middle heatmap), and negative regulators of triglyceride metabolism (lower heatmap). Genes labeled in magenta are members of diapause-development specialized paralog pairs. Specifically, enzymes related to triglyceride metabolism were strongly upregulated during diapause (left columns) and downregulated during development (right columns). Several positive and negative regulators of TG metabolism were also upregulated and downregulated in diapause respectively, though the pattern was more variable. (C) Comparison between triglyceride subclass levels between African turquoise killifish and red-striped killifish, shown as fold change in total lipid abundance. All triglycerides belonging to each class (PUFA, poly-unsaturated fatty acids cumulatively (PUFA) or with specifically with five (5) or six (6) unsaturated/double-bond sites respectively; Very Long-Chain FA, long-chain fatty acids that contain 22 carbons) were summed for this analysis (see Methods). The same developmental stage (pre-diapause stage) was compared between the two species. African turquoise killifish has a higher TG content at the Pre-Diapause stage compared to red-striped killifish. (D) Heatmap representing the fold change of all significant lipids species between diapause vs. development in the African turquoise killifish (left panels) and between the African turquoise killifish vs. red-striped killifish (development only, rightmost panel). Fold change values are plotted between each pair-wise comparison between diapause and development time points, or the two development time points. Lipids were included if significance was reached in any single comparison. The rightmost panel shows the fold change values of the same lipids in the African turquoise killifish compared to the red-striped killifish.

#### Figure S16

**A** Distribution of genes upregulated in diapause in paralogs and singletons

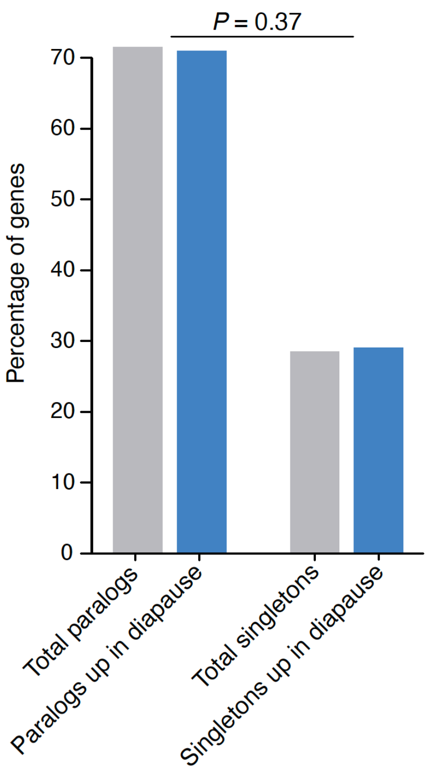


**B** Motif enrichment comparison for singletons and paralogs

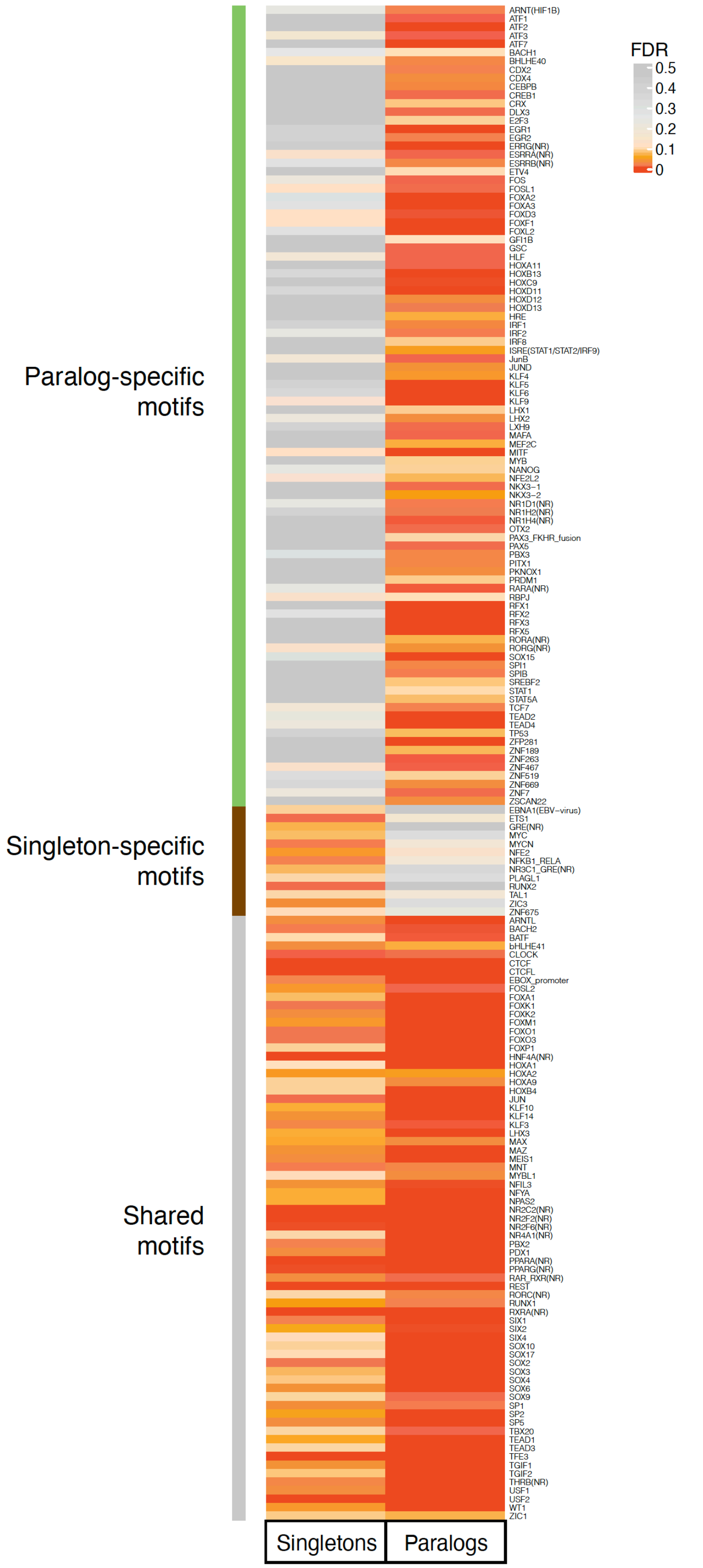

**Figure S16. Comparison of paralogs and singleton genes in the African turquoise killifish.** (A) Comparison of paralogs and singleton genes upregulated in diapause with their respective genome-wide expectation. Neither paralogs nor singletons are overrepresented among diapause-upregulated genes (*P* = 0.37 from Chi-squared test). (B) Transcription-factor binding motif enrichment in diapause-accessible chromatin regions (ATAC-seq peak) near singletons (left column) and paralogs (right column) in the African turquoise killifish. The majority of enriched motifs are either specific to paralogs or shared between paralogs and singletons with a minority being singleton specific (13 motifs), suggesting that the genomic regulatory landscape might be different for paralogs and singletons.

Figure S17

**A** Alignment-independent promoter motif enrichment between the African turquoise killifish and the lyretail killifish

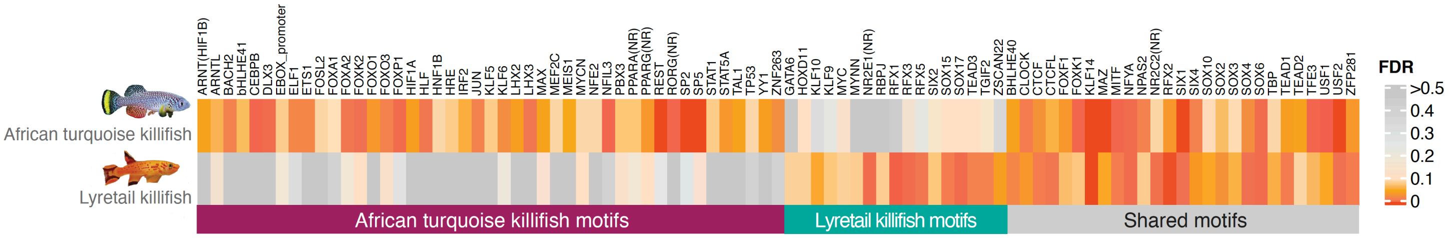


**B** Alignment-independent promoter motif enrichment between the African turquise killifish and the red-striped killifish

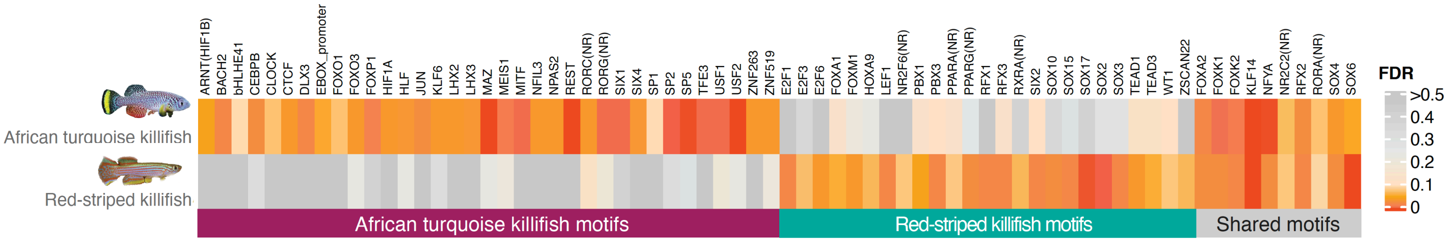

**Figure S17. Transcription-factor binding motifs enrichment using alignment-free approaches.** (A-B) Comparison of enriched transcription-factor binding motifs between African turquoise killifish promoters and lyretail killifish promoters (A) and red-striped killifish promoters (B). To produce an enrichment set without the use of a species whole genome multi-alignment, all accessible chromatin regions (ATAC-seq peaks) located in the promoters of diapause-specialized genes (African turquoise killifish) and their orthologs in lyretail or red-striped killifish were used. Similar to the alignment-based comparison (Fig. 4B, fig. S11), most promoter motifs are also species specific suggesting they evolved recently. Diapause specific motifs such as FOXO and REST are also only enriched in the African turquoise killifish promoters and not in lyretail or red-striped killifish promoters.

#### SUPPLEMENTARY TABLE

**Table S1**: Killifish and outgroup species used in this study.

| Common name | Scientific name | Group | Genome assembly |
|---|---|---|---|
| African turquoise killifish | *Nothobranchius furzeri* | African (with diapause) | Nfu_20140520 (*8*) |
| Lyretail killifish | *Aphyosemion australe* | African (without diapause) | MPIBA_Aaus_1.0 (*26*) |
| Red-striped killifish | *Aphyosemion striatum* | African (without diapause) | MPIBA_Aaus_1.0 (*26*) |
| South American killifish | *Austrofundulus limnaeus* | South American (with diapause) | Austrofundulus_limnaeus -1.0 (*9*) |
| Medaka | *Oryzias latipes* | Outgroup (without diapause) | ASM223467v1 (*130*) |
| Zebrafish | *Danio rerio* | Outgroup (without diapause) | GRCz11 (*131*) |

#### LIST OF SUPPLEMENTARY DATA FILES

**Data File S1:** Accession numbers and details of the datasets generated and used in this study.

**Data File S2:** Specialized paralogs in African turquoise killifish generated by OrthoFinder with 71 species used for the analysis.

**Data File S3:** Diapause-specific ATAC-seq peaks in African turquoise killifish, time of their origin, their closest genes, and their positive selection status.

**Data File S4:** Protein coding genes under positive selection in the ancestor of African turquoise killifish at the time of diapause evolution.

**Data File S5:** Enriched Gene Ontology functions for diapause gene expression of paralogs and diapause specific ATAC-seq data.

**Data File S6:** Enriched Gene Ontology functions for diapause specific ATAC-seq peaks under positive selection.

**Data File S7:** Upstream regulators of paralog gene expression predicted using Ingenuity Pathway Analysis (IPA).

**Data File S8:** Diapause-specific changes in the lipidome of the African turquoise and red-striped killifish.
